# Extracellular vesicles adhere to cells primarily by interactions of integrins and GM1 with laminin

**DOI:** 10.1101/2024.04.11.589011

**Authors:** Tatsuki Isogai, Koichiro M. Hirosawa, Miki Kanno, Ayano Sho, Rinshi S. Kasai, Naoko Komura, Hiromune Ando, Keiko Furukawa, Yuhsuke Ohmi, Koichi Furukawa, Yasunari Yokota, Kenichi G. N. Suzuki

## Abstract

Tumor-derived extracellular vesicles (EVs) have attracted significant attention, yet the molecular mechanisms that govern their specific binding to recipient cells remain elusive. Our in vitro study utilizing single-particle tracking demonstrated that integrin heterodimers comprising α6β4 and α6β1 and ganglioside, GM1 are responsible for the binding of small-EV (sEV) subtypes to laminin. EVs derived from four distinct tumor cell lines, regardless of size, exhibited high binding affinities for laminin but not for fibronectin, although fibronectin receptors are abundant in EVs and have functional roles in EV-secreting cells. Our findings revealed that integrins in EVs bind to laminin via the conventional molecular interface, facilitated by CD151 rather than by inside-out signaling of talin-1 and kindlin-2. Super-resolution movie observation revealed that sEV integrins bind only to laminin on living recipient cells. Furthermore, sEVs bound to HUVEC and induced cell branching morphogenesis in a laminin-dependent manner. Thus, we demonstrated that EVs predominantly bind to laminin on recipient cells, which is indispensable for cell responses.

**Summary:** Quantitative assessments using single-molecule imaging and super-resolution microscopy revealed that all extracellular vesicle subtypes derived from four distinct tumor cell lines, regardless of size, bind to laminin predominantly via CD151-facilitated integrin heterodimers and GM1, leading to response of recipient cells.

## Introduction

Extracellular vesicles (EVs) of a variety of sizes (∼30-1000 nm) contain a diverse array of cargo, including microRNAs, metabolites, and proteins, and play a critical role in intercellular communication (Valadi et al., 2007; Raposo and Stoorvogel, 2013; van Niel et al., 2018). EVs have attracted extensive attention in a wide range of research fields, and the number of published papers has increased dramatically (Théry et al., 2018; Witwer et al., 2021). In particular, several reports have suggested that tumor-derived EVs can induce phenotypic changes in recipient cells, creating a premetastatic niche that promotes cancer metastasis (Peinado et al., 2012; Ono et al., 2014). sEV studies might elucidate the underlying mechanism of the ‘seed and soil’ hypothesis (Paget, 1989), which posits that cancer cells tend to metastasize and proliferate in organs suitable for these activities (Hart and Fidler, 1980; Peinado et al., 2017). A previous proteomic study of sEVs revealed that integrin α6β4 and α6β1 in sEVs from 4175-LuT human breast cancer cells were associated with lung metastasis, while αVβ5 in small EVs (sEVs) from BxPC3 human pancreatic adenocarcinoma cells was linked to liver metastasis (Hoshino et al., 2015). In addition, several studies have shown that blocking integrin in sEVs through anti-integrin antibodies or integrin ligands inhibits the internalization of integrin by recipient cells (Nazarenko et al., 2010; Altei et al., 2020). However, no direct evidence that integrin heterodimers in sEVs bind to the extracellular matrix (ECM) components on recipient cell plasma membranes (PMs) has been reported. In addition, the binding affinities between EVs and diverse ECM components have yet to be determined via comparative analysis. Furthermore, the detailed molecular-level mechanisms that govern the selective binding of sEVs to recipient cell PMs remain elusive. Elucidating these mechanisms is critical for developing effective strategies to inhibit cancer metastasis (Möller and Lobb, 2020; Marar et al., 2021). Moreover, it is essential to validate the biological functions of sEVs and their roles in physiological and pathological processes (Gurung et al., 2021).

In this study, we aimed to investigate whether integrin heterodimers play a critical role in the binding of sEVs to ECM components on recipient cell PMs. Furthermore, we sought to elucidate the mechanisms of selective binding using our cutting-edge imaging techniques. Since we demonstrated that sEVs can be classified into several subtypes with different tetraspanin marker proteins (CD63, CD81, and CD9) and that each subtype has unique properties (Hirosawa et al., 2024, *Preprint*), it was crucial to examine the binding of each sEV subtype to ECM components. To this end, we first established an in vitro assay system that enabled us to quantitatively evaluate the binding of three sizes of EVs (small EVs, sEVs; medium-sized EVs, mEVs; and microvesicles; MVs) to ECM components on glass by single-particle tracking with single-molecule detection sensitivity. Second, we developed a state-of-the-art imaging system that enables the acquisition of super-resolution (dSTORM) movies, as opposed to static images. These advances enabled us to examine the binding affinities of various EV subtypes to ECM components on living cell PMs.

## Results

### Characterization of sEVs by single-particle tracking with single-molecule detection sensitivity

To investigate whether integrin subunits in tumor-derived sEVs are essential for selective binding to ECM components, we first characterized sEVs isolated from human prostate cancer (PC3) cells using an ultracentrifugation-based (pellet-down) method (Fig. S1A). These sEVs displayed circular morphology, with a diameter of 83 ± 19 nm (mean ± S.D.) (Fig. S1B), as determined by transmission electron microscopy (TEM) with negative staining. Consistently, the diameter measured by a tunable resistive pulse sensing instrument (qNano) was 69 ± 17 nm (mean ± S.D.) (Fig. S1C).

Recently, we demonstrated that PC3 cell-derived sEVs comprise several distinct subtypes, each harboring different tetraspanin marker proteins, as determined by single-particle imaging analysis (Hirosawa et al., 2024, *Preprint*). Approximately 40% of the sEVs contained the CD63, CD81, and CD9 tetraspanin marker proteins, while the remaining 60% of the sEVs contained only CD63. Furthermore, sEVs that contained only CD63 colocalized with caveolae in the recipient cell PMs, whereas those containing all three tetraspanin proteins did not. These findings indicate that the sEV subtypes interact differently with recipient cell PM structures (Hirosawa et al., 2024, *Preprint*). Therefore, in this study, to investigate the roles of integrin heterodimers in different sEV subtypes, we isolated sEVs from PC3 cells that stably expressed CD63-Halo7, CD81-Halo7, or CD9-Halo7. The tetraspanin-Halo7 in these sEVs was labeled by conjugation of the SaraFlour650T (SF650T) ligand, henceforth referred to as “sEV-tetraspanin-Halo7-SF650T”. We successfully observed single sEV-tetraspanin-Halo7-SF650T particles using total internal reflection fluorescence microscopy (TIRFM) with no contamination from free dye molecules or autofluorescence from non-labeled sEVs (Fig. S1D). Since the estimated average numbers of CD63-Halo7, CD81-Halo7, and CD9-Halo7 molecules per sEV particle are between 4.2 and 4.8 (Hirosawa et al., 2024, *Preprint*), according to the Poisson distribution, more than 98% of sEV particles should contain at least one Halo7-tagged tetraspanin protein. To standardize the concentrations of sEVs derived from different cell lines, we utilized single-particle imaging with single-molecule detection sensitivity to directly measure the number of sEV-tetraspanin-Halo7-SF650T particles bound to the antibody-coated glass. After incubating the sEV solution (2×10^10^ particles/ml) on antibody-coated glass at a dilution factor (df) of 1, 3 or 9, followed by three washes with HBSS, we obtained TIRFM images of single sEV-CD63-Halo7-SF650T particles (Fig. S1E). The number of sEVs bound to the glass decreased in accordance with the dilution factor (df), which allowed us to obtain a calibration curve (Fig. S1, F-J). By directly determining the concentration of sEV-tetraspanin-Halo7-SF650T and adjusting the concentration of the sEV solution to 0.8-4×10^10^ particles/ml, we were able to compare the binding numbers of sEVs derived from different cell types to ECM-coated glass.

### Knockout of integrin subunits

We employed CRISPR‒Cas9 gene editing to knock out the integrin β1, β4, α2 or α6 subunit in PC3 cells to investigate the role of these subunits in the binding of sEVs to ECM components. Western blotting analysis of the sEVs obtained from intact cells revealed the presence of the integrin β1, β4, α2, α6, α3, α5, and αV subunits, as well as greater abundances of the tetraspanin marker proteins CD63, CD81, and CD9 in comparison to those in parental cells (Fig. 1A). We confirmed that the integrin β1, β4, α2, and α6 subunits were indeed eliminated in sEVs derived from integrin-KO cells. Additionally, integrin α2 and α3 were below the detection limit in sEVs from integrin β1-KO cells (Fig. 1A). Moreover, integrin β4 was not detected in sEVs from integrin α6 KO cells, and integrin α6 expression was decreased in sEVs from β4 KO cells (Fig. 1A). Thus, the knockout of integrin subunits in sEVs often results in decreased expression levels of integrin heterodimer counterparts (Fig. 1 B) (Hynes, 2002), while the expression levels of other integrin subunits remain unaffected.

**Fig. 1.**
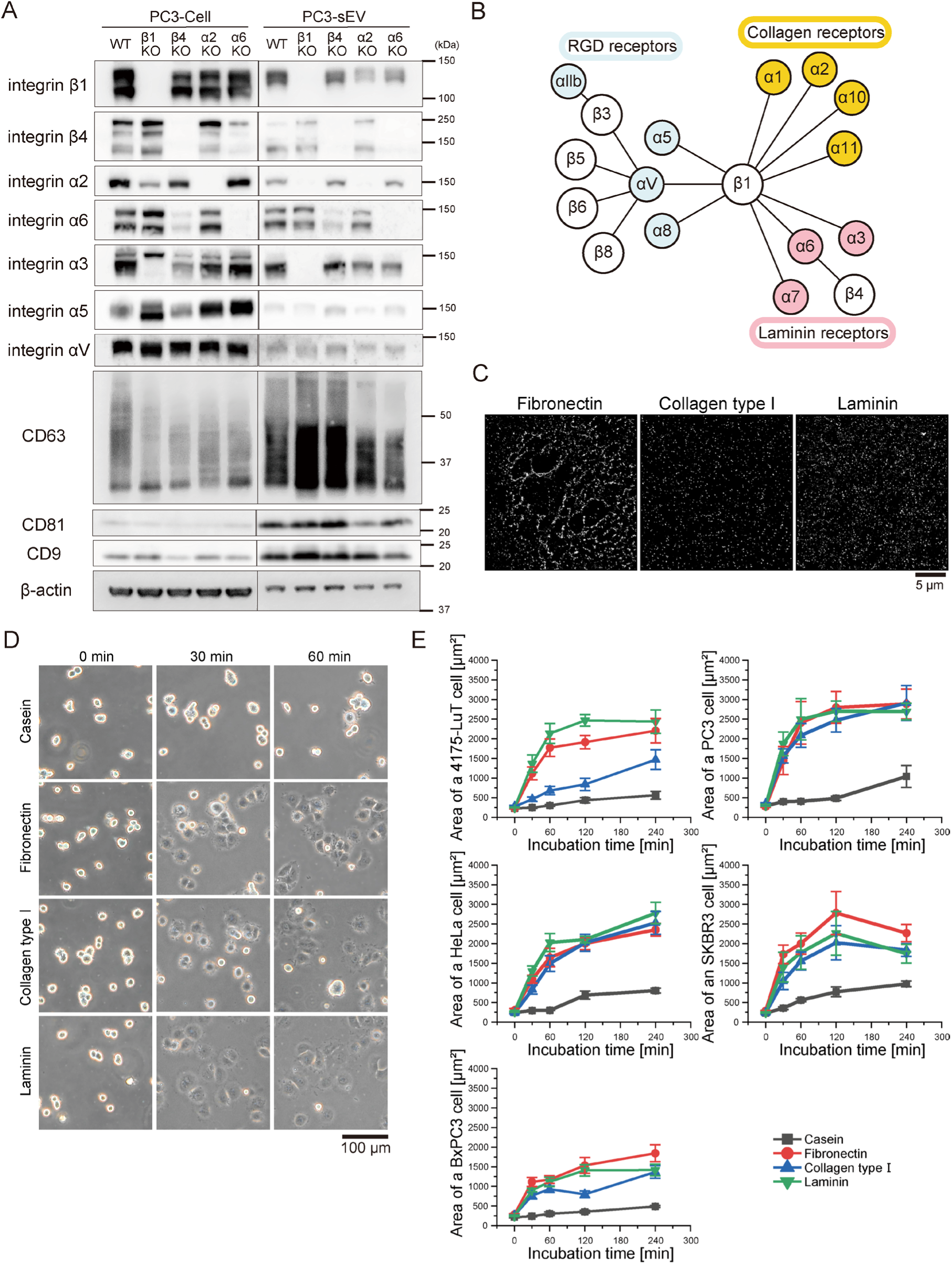
Western blotting of integrin subunits in PC3 cells and sEVs, and spreading assay of the cells that secreted sEVs in this study on all three ECM (fibronectin, laminin, and collagen type I) coated on glass. (A) sEVs were isolated from the cell culture supernatant of intact PC3 cells or from cells in which an integrin subunit (β1, β4, α2, α6) was knocked out via the CRISPR-Cas9 method. (B) The correlation map of integrin heterodimers and the ECM. (C) dSTORM images of the ECM (left-top: fibronectin, left-bottom: collagen type Ⅰ, right: laminin) coated on glass. (D) Images of HeLa cells attached to glass coated with ECM components (fibronectin, collagen type Ⅰ, laminin) or casein after 0, 30, or 60 min of incubation. (E) The time course of the area of five tumor cell lines on glass coated with ECM components (fibronectin, collagen type Ⅰ, laminin) or casein (*n* = 8 cells). Data are presented as the mean ± SE.

### Integrin β1 in tumor-derived sEVs plays a critical role in binding to laminin and collagen type *Ⅰ on glass*

Integrins are responsible for bidirectional signaling, namely, inside-out and outside-in signaling. For instance, intracellular signaling molecules such as talin can activate integrins (Hynes, 2002; Moser et al., 2009; Lu et al., 2022). Although sEVs contain integrin heterodimers of ECM receptors, this does not necessarily imply that the integrin heterodimers in sEVs can bind to the ECM. Hence, to investigate whether integrin subunits in tumor-derived sEVs are responsible for sEV binding to ECM components, we first performed in vitro binding assays. dSTORM images of immunostained fibronectin, laminin, and collagen type I coated on glass showed that all the ECM components were attached to and evenly dispersed on the glass surfaces (Fig. 1C). Furthermore, almost all the cells that secreted sEVs in this study attached to and spread on all three ECM components coated on glass at similar velocities (Fig. 1, D and E), indicating that integrin heterodimers that act as receptors for all three ECM components are active on the cell PMs.

Fig. 2A shows TIRFM images of individual sEV-CD63-Halo7-SF650T particles derived from wild-type PC3 cells (PC3-WT) and integrin β1 KO cells (PC3-β1KO) adhered to glass surfaces coated with fibronectin, laminin, or collagen type I. Quantitative analysis revealed that the number of all subtypes of sEV-tetraspanin-Halo7-SF650T particles bound to fibronectin per image area (82 μm×82 μm) was very small; therefore, no reduction in quantity due to integrin β1 KO was observed (left in Fig. 2B, Fig. S2, A and B, and Table S1). Conversely, the numbers of all subtypes of sEV-tetraspanin-Halo7-SF650T particles attached to collagen type I were higher than those bound to fibronectin and significantly decreased by integrin β1 KO (middle in Fig. 2B, Fig. S2, A and B, and Table S1). Moreover, all subtypes of sEV-tetraspanin-Halo7-SF650T particles bound to laminin in considerably higher numbers than they bound to fibronectin and collagen type I, and the number decreased drastically due to integrin β1 KO (right in Fig. 2B, Fig. S2, A and B, and Table S1). These findings explicitly reveal that integrin β1 in PC3-derived sEVs is responsible for the binding of sEVs to laminin and collagen type Ⅰ.

**Fig. 2.**
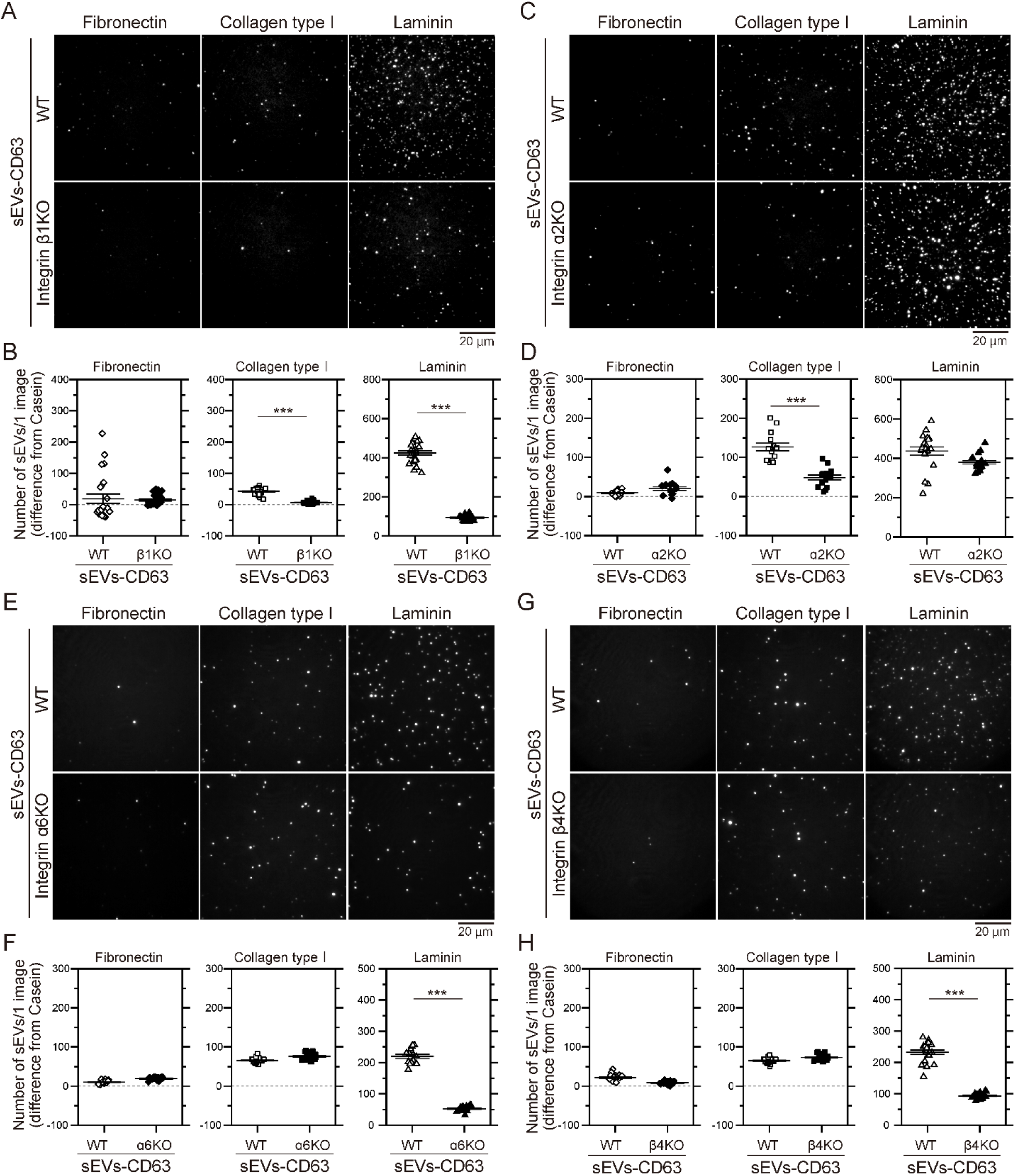
Integrin α2β1 in sEVs derived from PC3 cells is responsible for the binding of CD63-containing sEVs to collagen type I, and integrins α6β1 and α6β4 are responsible for the binding to laminin. (A, C, E, and G) Single-particle fluorescence images of sEV-CD63Halo7-SF650T on glass coated with fibronectin, collagen type I, and laminin before and after the KO of integrin β1 (A), integrin α2 (C), integrin α6 (E), and integrin β4 (G). (B, D, F, and H) The numbers of sEVs attached to glass coated with these extracellular matrix molecules before and after the KO of integrin β1 (B), integrin α2 (D), integrin α6 (F), and integrin β4 (H). Data are presented as the mean ± SE. n.s., non-significant difference; \**P*<0.05; \*\**P*<0.01; \*\*\**P*<0.001 according to Welch’s t-test (two-sided).

The surface of sEVs derived from myeloma cells and isolated by ultracentrifugation is covered by fibronectin, which serves as a heparan sulfate-binding ligand. Consequently, the enrichment of fibronectin in sEVs marginally (by a factor of approximately 1.2) amplifies the interactions between sEVs and recipient cells (Purushothaman et al., 2016). Therefore, we performed immunoprecipitation experiments to examine the presence of fibronectin and/or laminin on the surface of sEVs derived from PC3 cells (Fig. S3A). However, neither fibronectin nor laminin was detected in the fraction isolated by the immunoprecipitation of sEVs with the anti-CD63 antibody (Fig. S3B). Furthermore, we did not detect tetraspanin marker proteins (CD63, CD81, or CD9) in the fraction obtained through the immunoprecipitation of sEVs using anti-fibronectin antibody (Fig. S3C). These results unequivocally showed that neither laminin nor fibronectin was present on the surface of PC3 cell-derived sEVs.

### Integrin α2 in tumor-derived sEVs is responsible for binding to collagen type Ⅰ on glass

The in vitro assay revealed that in all the examined sEV subtypes, integrin β1 was critical for binding to collagen type I and laminin on glass. Accordingly, we focused on the function of integrin heterodimers containing the β1 subunit. Since the integrin α2β1 heterodimer is a receptor for collagen type I, we examined the role of the integrin α2 subunit. Fig. 2C shows TIRFM images of individual particles of sEV-CD63Halo7-SF650T derived from PC3-WT and PC3-α2KO cells, which attached to ECM components on glass surfaces. Notably, the expression levels of other integrin subunits in PC3-α2KO cells were comparable to those in PC3-WT cells (Fig. 1A). Quantitative analysis revealed a significant decrease in the number of all subtypes of sEV-tetraspanin-Halo7-SF650T particles derived from α2KO cells bound to collagen type I, while the number that bound to laminin was not significantly reduced (Fig. 2D, Fig. S2, C and D, and Table S1). These findings, in conjunction with the results of β1KO, unequivocally show the crucial involvement of integrin α2β1 heterodimers in the binding of sEVs to collagen type Ⅰ.

### Integrins α6 and β4 in tumor-derived sEVs are responsible for binding to laminin on glass

We next investigated the involvement of the counterpart in β1-containing integrin heterodimers in the binding of sEVs to laminin. As integrin α6β1 is a receptor for laminin, we aimed to determine whether the absence of integrin α6 affects the binding of sEVs to laminin on glass. To this end, we performed a western blotting experiment and detected a lack of the integrin β4 subunit in sEVs derived from PC3-α6KO cells. In addition, we observed a considerable decrease in the expression level of the integrin α6 subunit in EVs derived from β4KO cells (Fig. 1A). Therefore, it is plausible that the expression levels of integrins α6 and β4 in sEVs are closely linked. Fig. 2, E and G show TIRFM images of single particles of sEV-CD63Halo7-SF650T derived from PC3-WT, PC3-α6KO, and PC3-β4KO cells, which were attached to ECM components on glass. We discovered that knocking out integrin α6 or integrin β4 considerably impaired the binding of all subtypes of sEV-tetraspanin-SF650T to laminin but did not significantly affect the binding to fibronectin or collagen type I (Fig. 2, F and H, Fig. S2, E-H, and Table S1). These findings explicitly show that integrin heterodimers α6β1 and α6β4 in sEVs are responsible for binding to laminin on glass.

Thus, single-particle imaging of PC3 cell-derived sEVs on glass revealed that integrin α2β1 in sEVs was critical for binding to collagen type Ⅰ, while integrin α6β1 and integrin α6β4 in sEVs were responsible for binding to laminin (Fig. 2, Fig. S2, and Table S1). Interestingly, only small amounts of all the PC3 cell-derived sEVs bound to fibronectin on glass despite the presence of fibronectin receptors such as integrin α5β1, αVβ1 and αVβ5 in the PC3-derived sEVs (Fig. 1, A and B). Notably, these fibronectin receptors are functional in PC3 cells, as evidenced by their comparable spread on glass surfaces coated with laminin, fibronectin, or collagen type I (Fig. 1E).

### GM1, but not other gangliosides, in sEVs specifically binds to laminin

Our results showed that the knockout of integrin α6, β4, or β1 greatly reduced but did not eliminate the binding affinity of PC3-derived sEVs for laminin, suggesting that other molecules may be involved in the binding. Previous reports have suggested that laminin may bind directly to GM1 on dorsal root ganglion neurons (Ichikawa et al., 2009). Since our dot blot experiments showed that the relative density of GM1 (Fig. S4A) in sEVs derived from PC3 cells was approximately 29.7 ± 8.1 (mean ± S.E., n=3) times greater than that in PC3 cells (Fig. S4B), GM1 might play a role in the binding of sEVs to laminin. To investigate whether GM1 in sEVs binds directly to laminin, we sought to isolate cell lines that abundantly expressed only one type of ganglioside by overexpressing ganglioside synthase in cells to produce or deplete gangliosides, as reported previously (Yamashiro et al., 1995; Yesmin et al, 2023). However, this strategy did not work in PC3 cells because the amounts of gangliosides produced or depleted by the synthase returned to the original levels during cell culture. Therefore, we applied this method instead to B78 cells, a subclone of B16 melanoma cells that can predominantly express one type of ganglioside (Fig. S5). The number of sEVs that bound to laminin, normalized by the ratio of the expressed amount of integrin α6/CD81 in the sEVs (Fig. S4C), was approximately 3-fold greater for sEVs derived from B78 cells predominantly expressing GM1 than for sEVs derived from B78 cells expressing GM2, GM3, GD2 and GD2/GD3 (Fig. S4, A, D and E). Regardless of which gangliosides were expressed, the number of B78 cell-derived sEVs bound to fibronectin was low (Fig. S4, F and G). These results suggest that GM1, but not the other examined gangliosides in sEVs, might bind to laminin and that all the gangliosides in sEVs might not bind to fibronectin.

To further examine whether GM1 binds directly to laminin, we investigated whether liposomes of ∼100 nm in diameter containing GM1 bound to laminin but not to fibronectin. We used liposomes containing 1.0 mol% GM1, which was estimated to be equivalent to the concentration in sEVs for the binding assay (also see the Method section). Fig. S4, H and I demonstrate that liposomes containing GM1 bound extensively to laminin, while those containing other gangliosides (GM2, GM3, GD1a, GD2, and GD3) showed very little binding to laminin. Conversely, liposomes hardly bound to fibronectin regardless of which gangliosides they contained (Fig. S4, H and J). These results unequivocally establish that GM1 in sEVs directly binds to laminin but not to fibronectin.

We subsequently assessed the relative contributions of GM1 and integrin β1 in PC3-WT cell-derived sEVs to laminin binding by competitive binding inhibition assay. The number of sEVs-CD63Halo7-TMR derived from PC3-WT cells bound to laminin-coated glass decreased by 61% after integrin β1 KO (Fig. S4, K and L). Furthermore, the numbers of the sEVs from PC3-WT and integrin β1 KO cells bound to laminin were reduced by 18% and 34%, respectively, in the presence of a high concentration (0.5 mM) of GM1 glycan (Fig. S4, K and L). These results show that while integrin heterodimers in the sEVs predominantly mediate laminin binding, GM1 glycan in the sEVs also partially facilitates the interaction of the sEVs with laminin.

### All the sEVs, mEVs and MVs derived from four tumor cell lines exhibited marked binding affinity for laminin but minimal binding to fibronectin

To investigate whether the high binding affinity for laminin and the low binding affinity for fibronectin are general features of tumor-derived sEVs, we conducted in vitro binding assays of sEV-CD63-Halo7-SF650T particles derived from four other types of tumor cells. These sEVs were purified by ultracentrifugation (200,000×g for 4 h). The diameters of the sEVs derived from 4175-LuT, BxPC3, HeLa and SKBR3 cells, which were assessed via qNano, were 83 ± 16 nm, 97 ± 9 nm, 83 ± 12 nm and 75 ± 8 nm (mean ± S.D.), respectively (Fig. 3A). Quantitative analysis of the TIRFM images revealed that the numbers of sEV particles derived from four tumor cell lines (4175-LuT, PC3, BxPC3, and SKBR3) that bound to laminin on glass were considerably greater than those that bound to fibronectin on glass (Fig. 3, B and C). On the other hand, the numbers of sEVs derived from HeLa cells that bound to either fibronectin or laminin were very low (Fig. 3C). These results explicitly demonstrated that tumor-derived sEVs primarily bind to laminin on glass and exhibit low affinity for fibronectin.

**Fig. 3.**
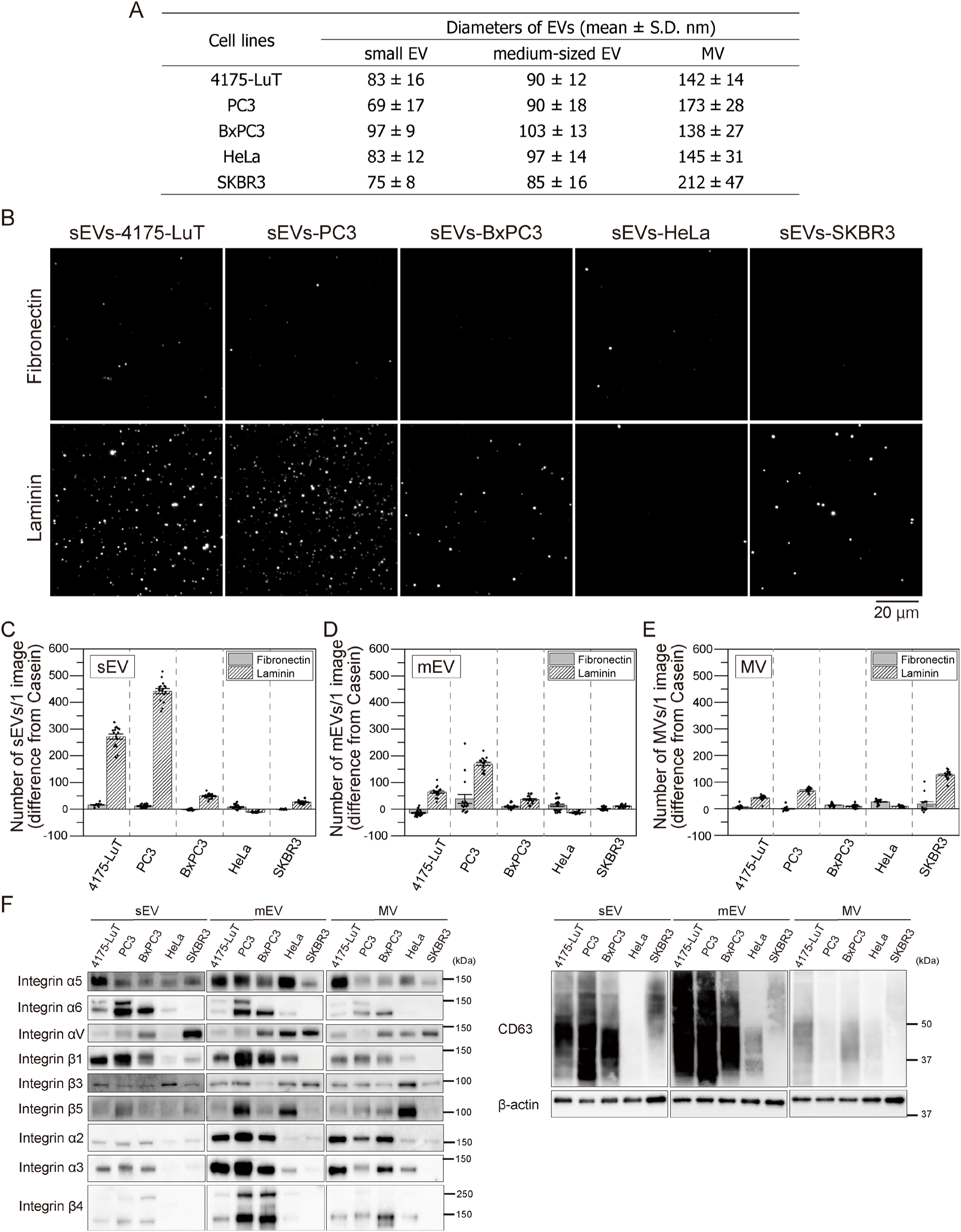
Tumor-derived sEVs bind much more strongly to laminin than to fibronectin. (A) The diameters of sEVs, mEVs, and MVs derived from the 4175-LuT, PC3, BxPC3, HeLa, and SKBR3 cell lines were measured by qNano. (B) Single-particle fluorescence images of sEV-CD63Halo7-SF650T particles derived from the 4175-LuT, PC3, BxPC3, HeLa, and SKBR3 cell lines on glass coated with fibronectin or laminin. (C-E) The numbers of EV-CD63Halo7-SF650T particles (sEVs (C), mEVs (D), and MVs (E)) attached to glass coated with ECM components. Data are presented as the mean ± SE. The numbers of sEVs, mEVs, and MVs derived from all cell lines applied to laminin-or fibronectin-coated glass were adjusted to the same. (F) Western blotting of integrin subunits in sEVs, mEVs, and MVs derived from 4175-LuT, PC3, BxPC3, HeLa, and SKBR3 cells.

The diameter of EVs isolated by ultracentrifugation depends on factors such as the sample volume, centrifugation tube, centrifugal acceleration, and centrifugal duration (Théry et al., 2018). Therefore, to investigate whether the stronger binding affinity of EVs for laminin is a general phenomenon, we quantified the number of larger EVs bound to ECM components coated on glass. This experiment was necessary because sEVs and MVs are reported to contain distinct membrane components (Jeppesen et al., 2019). To achieve this goal, we isolated larger EVs by ultracentrifugation at slower rates. The mEVs were isolated by low-speed centrifugation (50,000×g for 30 min) and had diameters of 90 ± 12 nm (4175-LuT), 90 ± 18 nm (PC3), 103 ± 13 nm (BxPC3), 97 ± 14 nm (HeLa), and 85 ± 16 nm (SKBR3) (Fig. 3A). Our in vitro binding assay showed that these mEVs share similar characteristics with sEVs (Fig. 3D). Western blotting (Fig. 3F) revealed that fibronectin receptors, such as integrin α5β1 heterodimers, were expressed in the EVs derived from all tested types of tumor cells. In addition, the diameters of the MVs purified by lower-speed centrifugation (10,000×g for 30 min) were 142 ± 14 (4175-LuT), 173 ± 28 (PC3), 138 ± 27 (BxPC3), 145 ± 31 (HeLa), and 212 ± 47 (SKBR3) (Fig. 3A). Since CD63 was not highly concentrated in MVs (Fig. 3F), the MVs were stained with the lipid probe (ExoSparkler exosome membrane labeling kit-Deep Red, Dojindo), and the numbers of MVs bound to laminin and fibronectin were quantified by TIRFM (Fig. 3E). Our results indicate that MVs derived from 4175-LuT, PC3, and SKBR3 cells exhibit greater binding affinity to laminin than to fibronectin. However, the numbers of MVs derived from BxPC3 and HeLa cells that bound to laminin and fibronectin were very low (Fig. 3E). Thus, despite the presence of fibronectin receptors such as integrin α5β1 and αVβ3 in all sizes of EVs (Fig. 3F), these proteins exhibited minimal binding to fibronectin. Notably, tumor cells spread on fibronectin- and laminin-coated glass at comparable rates (Fig. 1E), indicating that all receptors for ECM components on the tumor cell PMs were functional. These results demonstrate that the laminin receptor is markedly active and the fibronectin receptor activity is low in EVs derived from four tumor cell lines (4175-LuT, PC3, BxPC3, and SKBR3).

### The integrin β1 and α6 subunits in sEVs are responsible for binding to cell plasma membranes

Although the in vitro assay showed that integrin heterodimers play critical roles in binding to laminin and collagen type I on glass (Fig. 2 and Fig. S2), it remains unknown whether these integrin heterodimers also mediate the binding of sEVs to the recipient cell PM. To address this question, we quantitatively determined the number of sEV-CD63-SF650T particles bound to the PM of the recipient cell. Immunofluorescence imaging revealed that while human fetal lung fibroblast (MRC-5) PMs were extensively coated with fibronectin, collagen type I, and laminin, these coatings were poor in human marrow stromal (HS-5) cells (Fig. 4, A and B). The laminin α5 isoform was significantly more abundant on MRC-5 PMs than the α1 isoform, indicating that laminin-511, the same isoform used in the in-vitro assay, represents the predominant isoform on MRC-5 PMs (Fig. 4A). Then, time course of sEVs bound to the basal PMs of recipient cells was observed by TIRFM. After 30 min of incubation, the number of sEV-CD63-SF650T particles derived from PC3-WT cells that bound to the basal PM of MRC-5 cells was substantially greater than the number of sEV-CD63-SF650T particles derived from PC3-β1KO cells (p = 0.046; two-sided Welch’s t-test) or PC3-α6KO cells (p = 0.0082) (Fig. 4, C, E, F and H). In contrast, the number of PC3-β1KO cell-derived sEVs that bound to HS-5 cell basal PMs was indistinguishable from that of PC3-WT cell-derived sEVs (p = 0.55) (Fig. 4, D and G). To investigate the binding of individual sEVs to both the apical and basal PMs of recipient cells, we employed confocal fluorescence microscopy after chemical fixation. The number of PC3-WT-derived sEVs bound to MRC-5 PMs after 60-min incubation was approximately twice greater than that of PC3-β1KO-derived sEVs (p = 0.0022), while the numbers of these sEVs bound to HS-5 cell PMs were minimal and statistically indistinguishable (p = 0.30) (Fig. 4I and Fig. 4J). The number of sEVs on MRC-5 PMs was approximately 11-fold greater than that on HS-5 cell PMs (Fig. 4J). These results explicitly indicate that integrin β1 and integrin α6 in sEVs are involved in binding to the MRC-5 PM of which surface is densely covered by ECM components.

**Fig. 4.**
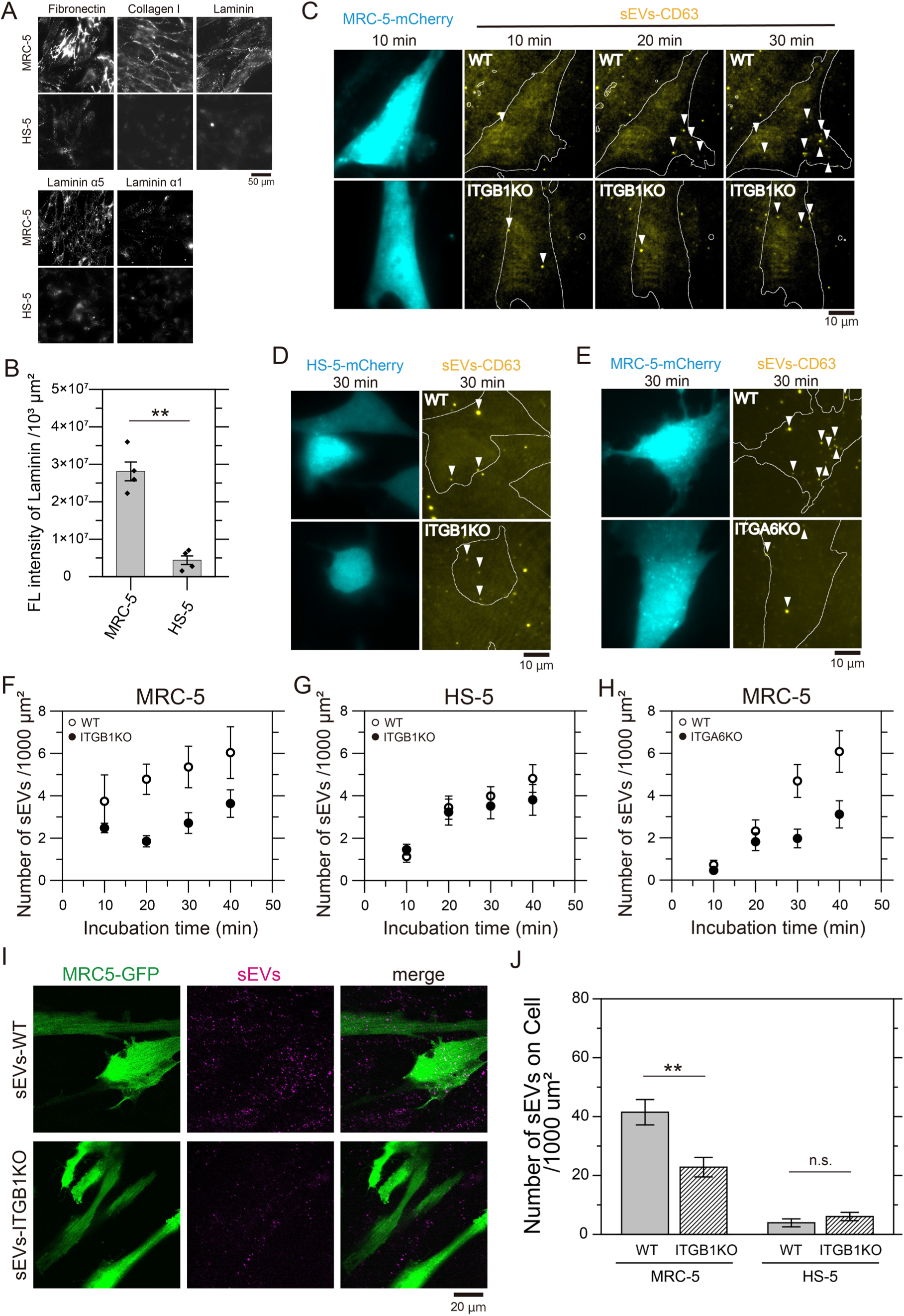
Integrins β1 and α6 in sEVs and the extracellular matrix (ECM) are responsible for the binding of sEVs to cell plasma membranes. (A) Immunofluorescence images of fibronectin, collagen type I, laminin, laminin α5, and laminin α1 in human normal embryonic lung fibroblast (MRC-5) cells and human bone marrow stromal (HS-5) cells. (B) The fluorescence intensities of laminin on cells in 1000 μm^2^. (C) Total internal reflection fluorescence (TIRF) images of MRC-5 cells expressing mCherry and sEV-CD63Halo7-SF650T particles (arrowhead) attached to cell membranes after 10, 20, and 30 min of incubation. sEVs isolated from the cell culture supernatant of intact PC3 cells (top panels) and from the cell culture supernatant of integrin β1 KO PC3 cells (bottom panels) bound to the basal surface of MRC-5 cells. (D) TIRF images of HS-5 cells and sEV-CD63Halo7-SF650T particles (arrowhead) attached to the basal surface of cell membranes after 30 min of incubation. sEVs were isolated from intact PC3 cells (top panels) and integrin β1 KO PC3 cells (bottom panels). (E) TIRF images of MRC-5 cells and the attached sEV-CD63Halo7-SF650T particles derived from intact PC3 cells (top panels) and from integrin α6 KO PC3 cells (bottom panels) after 30 min of incubation. (F-H) Time course of the numbers of intact sEVs and integrin β1-KO sEVs attached to the basal surface of MRC-5 cell membrane (*n* = 8 images) (F) and to the basal surface of HS-5 cell membrane (*n* = 15 images) (G) per 1000 μm^2^. (H) Time course of the numbers of intact sEVs and integrin α6 KO sEVs attached to the basal surface of MRC-5 cell membrane per 1000 mm^2^ (*n* = 12 images). Data are presented as the mean ± SE. (I) Fluorescence images of MRC-5 cells expressing GFP and sEV-CD63Halo7-TMR by confocal microscopy. sEVs isolated from the cell culture supernatant of wild-type (WT) PC3 cells (top panels) and from that of integrin β1 KO PC3 cells (bottom panels). The cells were fixed after treatment of the sEVs for 1 hour. While all the cells do not necessarily express GFP and many fluorescent spots of sEV-CD63Halo7-TMR are observed outside of GFP-expressing cells, we can quantify the number of sEV-CD63Halo7-TMR bound to both the apical and basal surface of the cell PM by counting the number of the fluorescent EV spots on cells expressing GFP. (J) The numbers of sEVs attached to MRC-5 and HS-5 cell PMs by confocal fluorescence microscopy. Data are presented as the mean ± SE. n.s., non-significant difference; \**P*<0.05; \*\**P*<0.01; \*\*\**P*<0.001 according to Welch’s t-test (two-sided).

### Integrins in sEV bind to laminin via the conventional molecular interface

We investigated whether integrin β1 in PC3-derived sEVs adopts an active conformation upon binding to laminin. To address this, we performed immunostaining of integrin β1 in sEV-CD63Halo7 bound to either uncoated glass or laminin-coated glass using antibodies specific for activated integrin β1 (clones HUTS-4 and HUTS-21, Luque et al., 1996) and a general anti-integrin β1 antibody (clone P5D2). Nearly all sEV-CD63Halo7-TMR particles bound to uncoated glass or laminin-coated glass were stained with the P5D2 antibody (Fig. 5, A-C). However, only 20-30% of the sEV particles on uncoated glass were stained with HUTS-4 and HUTS-21, whereas 60-70% of those bound to laminin-coated glass were positively stained with these activation-specific antibodies (Fig. 5, A-C). These results unequivocally show that integrin β1 in sEVs adopts an active conformation upon laminin binding.

**Fig. 5.**
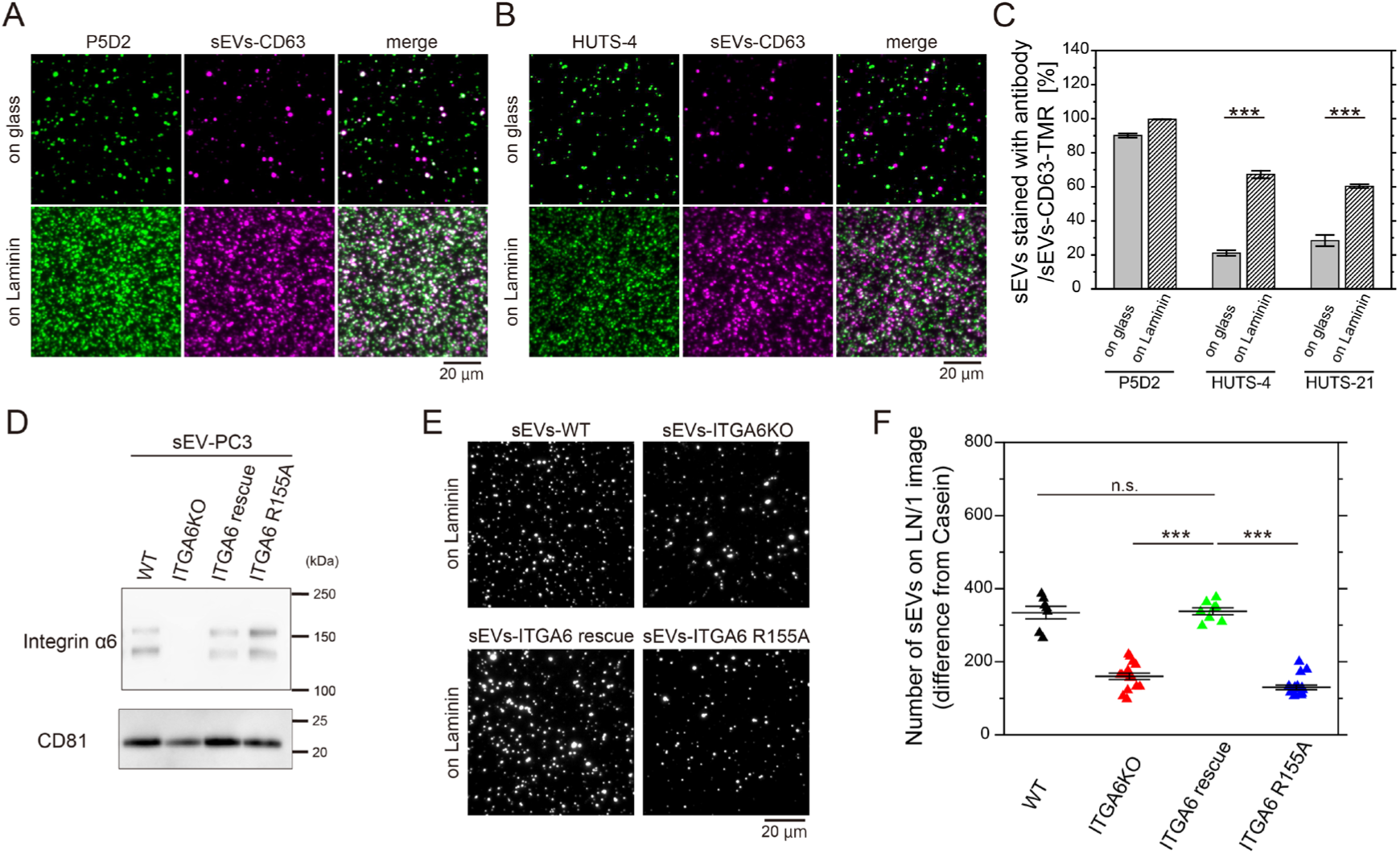
Integrins in sEV bind to laminin through conventional integrin-laminin interactions. (A and B) TIRF images of sEV-PC3-CD63Halo7-TMR stained with anti-activated integrin β1 HUTS-4 (A) and anti-integrin β1 P5D2 (B) on uncoated glass (top) or laminin-coated glass (bottom). (C) Colocalization ratio of fluorescent spots of sEVs stained with the antibody to those of sEV-PC3-CD63Halo7-TMR. (D) Western blot analysis of sEVs derived from wild-type, integrin α6 KO, integrin α6-rescued, and integrin α6 R155A-expressing PC3 cells. (E) Fluorescence images of the sEVs derived from wild-type, integrin α6 KO, integrin α6-rescued, and integrin α6 R155A-expressing PC3 cells bound to laminin on glass. (F) Quantification of sEVs bound to laminin on glass under the conditions described in (E). Data are presented as the mean ± SE. n.s., non-significant difference; \**P*<0.05; \*\**P*<0.01; \*\*\**P*<0.001 according to Welch’s t-test (two-sided).

Next, we investigated whether integrin α6 in sEVs binds to laminin via the same molecular interface as the conventional interaction in cells. To this end, we compared the laminin-binding capacity of sEVs derived from integrin α6 KO-PC3 cells expressing the R155A dominant-negative mutant of integrin α6 (Arimori et al., 2021) with that of sEVs derived from integrin α6-rescued cells. The levels of rescued integrin α6 and the R155A mutant in sEVs were comparable to the levels of endogenous integrin α6 in sEVs derived from PC3-WT cells (Fig. 5D). The number of sEVs containing the integrin α6 R155A mutant bound to laminin was significantly lower than those containing endogenous or rescued wild-type integrin α6 but was comparable to that of sEVs from integrin α6 KO PC3 cells (Fig. 5, E and F). These results explicitly show that the interaction between integrin heterodimers in sEVs and laminin relies on the same molecular interface as the conventional interaction in cells.

### Pseudo real-time super-resolution movie observation demonstrated that sEVs bind predominantly to laminin on the living cell PM

Having demonstrated the binding of integrin α2β1 in PC3 cell-derived sEVs to collagen type I on glass and the binding of both integrin α6β1 and integrin α6β4 in sEVs to laminin on glass, we next examined whether these sEVs also bind to ECM components on the living recipient cell PM. To directly assess this binding, we performed simultaneous single-particle tracking of PC3 cell-derived sEVs-CD63-Halo7-TMR particles and super-resolution microscopy observation of ECM components on the living MRC-5 cell PM (Fig. 6, A-E). Since both sEVs and ECM components exhibited slow movement on the cell PM and their positions changed throughout the observation, we sought to more accurately analyze the colocalization between sEVs and ECM components by acquiring super-resolution “dSTORM movies” of ECM components instead of the still images. For this purpose, we acquired dSTORM images by the single-molecule observation of ECM components immunostained with SF650B-conjugated antibodies at high speed (200 frames/second) for 3504 frames. To obtain one dSTORM image, data acquisition was performed for 1002 frames and then repeated by shifting the initial frames backward by 6 frames, resulting in a total of 417 dSTORM images (Fig. 6A). By connecting these dSTORM image sequences, we generated a pseudo-real-time “dSTORM movie”. Images of individual sEV particles were concurrently recorded at 200 frames/second, and images averaged over 6 frames were connected to create the movie. The pseudo-real-time movies of the ECM components and sEV particles (33 frames/second) were superimposed (Fig. 6A).

**Fig. 6.**
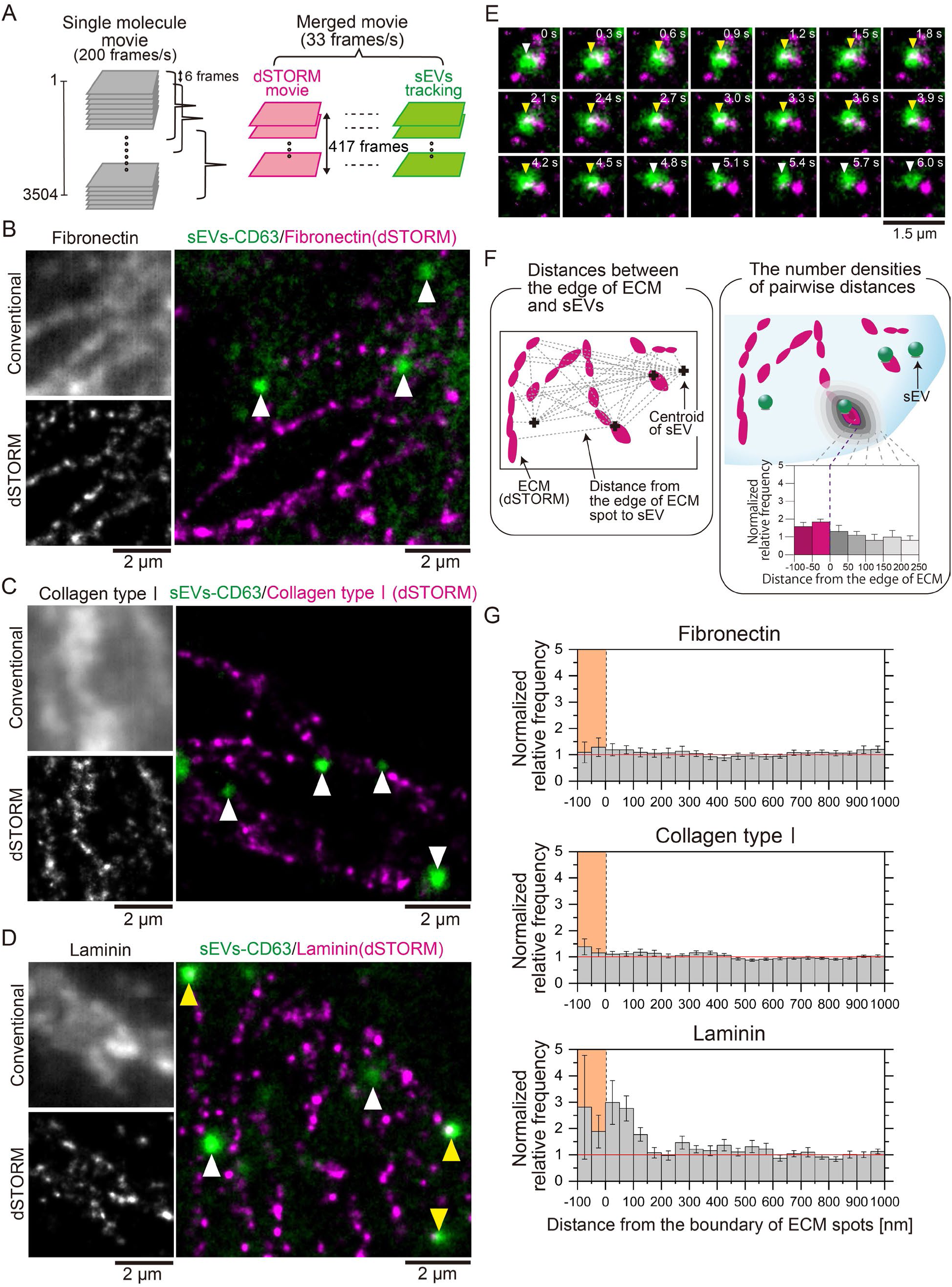
sEVs bind to laminin on the plasma membranes of living MRC-5 cells, as revealed by pseudo-real-time super-resolution movie observation. (A) Schematic diagram for generating merged movies of dSTORM images of the ECM structures and single-particle images of sEVs. Data acquisition was performed by observing single fluorescent molecules of SF650B at 200 frames/s (3,504 frames), and dSTORM images were reconstructed using the data acquired every 5.0 s (=1,002 frames). The first dSTORM image was reconstructed using frames 1-1,002 and the process was then repeated by shifting the initial frames backward by 6 frames; thus, 417 dSTORM images were obtained. These dSTORM still images were connected to construct the “dSTORM movie”. The single-particle movies of sEVs were subjected to a rolling-average for 6 frames and synchronously merged with the dSTORM movie. (B-D) Conventional immunofluorescence images (top-left) and dSTORM images (bottom-left) of the ECM structures (B: fibronectin, C: collagen type I, D: laminin) on the cells. dSTORM images of the ECM structures (magenta) and single-particle images of sEV-CD63Halo7-TMR particles (green) were merged (right). sEVs localized near (<100 nm) the boundary of ECM structures and sEVs localized alone are indicated by yellow and white arrowheads, respectively. (E) The image sequence (every 0.3 sec) of laminin obtained by dSTORM (magenta) and a single sEV-CD63Halo7-TMR particle (green) on the living MRC-5 cell membrane. sEVs colocalized with laminin, as indicated by the yellow arrowhead. (F) Colocalization analysis method. The normalized relative frequency was defined as the ratio of the average value of the spatial pair correlation function of the actual image to that of randomly distributed spots generated by a computer. Zero on the x-axis indicates the contour of the ECM structures in the dSTORM images determined by the kernel density estimation (KDE) method. When sEVs are enriched near the ECM structures, the normalized relative frequency is greater than 1. (G) Probability density analysis of the sEVs and ECM structures. The colored areas indicate regions within the ECM structures. The sEV-CD63Halo7-TMR particles localized near the contour of laminin (*n* = 20 cells).

Conventional immunofluorescence microscopy failed to visualize the fibrillary structure of ECM components on living cells (left-top panels of Fig. 6, B-D). In contrast, live-cell dSTORM movie observation enabled clear visualization of this structure (left-bottom panels of Fig. 6, B-D). We found that sEVs frequently colocalized with the structure of laminin on the cell PM, as indicated by the yellow arrowheads in Fig. 6D (right) (Video 3), and that the duration of colocalization occasionally persisted for longer than 4 seconds (Fig. 6E and Video 4). Moreover, we observed scarcely any colocalization events between the sEVs and the fibrillary structures of fibronectin (Fig. 6B, right, and Video 1) or collagen type Ⅰ (Fig. 6C, right, and Video 2). To quantitatively analyze the colocalization events, we measured the nearest distance from an edge of the ECM structure to a centroid of the sEV spot and performed this measurement for all pairs of ECM structures and sEV particles. The data were normalized using an in silico distribution of randomized sEV spots, and subsequently, we obtained histograms showing the distribution of the normalized relative frequency of sEVs at each distance from the edge of the ECM structures (Fig. 6F). Our analysis revealed that sEVs were significantly enriched near the edges of laminin structures at distances ranging from −100 to 150 nm on the PMs of living MRC-5 cells (Fig. 6G, bottom). In contrast, we did not observe the presence of sEVs near fibronectin or collagen type Ⅰ structures on the MRC-5 PM (Fig. 6G, top and middle, respectively). Our results indicate that PC3 cell-derived sEVs adhere predominantly to laminin on the recipient cell PM. Notably, we did not observe any binding of sEVs to collagen type I on the PM, although sEVs did bind to collagen I on glass (Fig. 2, A-D and Fig. S2, A-D). This difference may be attributed to the considerably lower binding of sEVs to collagen type I than to laminin, as demonstrated in the glass assay (Fig. 2 and Fig. S2).

### Talin-1 and kindlin-2 do not facilitate the binding of integrin heterodimers in EVs to laminin

Talin binds to the cytoplasmic tail of integrin β subunits and thereby disrupts the transmembrane association of integrin heterodimers, induces extended-open conformations, and promotes the binding of integrin heterodimers to all ECM components (Sun et al., 2019). Talin-1, but not talin-2, is an important factor for activating integrin β1 in prostate cancer (PCa) cells, which facilitates bone metastasis (Jin et al., 2015). The phosphorylation of talin-1 at Ser 425 promotes the activity of integrin β1, accelerating metastasis (Jin et al., 2015). Furthermore, Cdk5 kinase activity is responsible for talin-1 phosphorylation at Ser 425 (Huang et al., 2009), which subsequently triggers integrin β1 activation (Jin et al., 2015). Moreover, the present study showed that the presence of fibronectin receptors in EVs does not correspond to binding with fibronectin. Therefore, we hypothesized that factors other than the presence of ECM receptors are involved in regulating the binding of EVs with ECM components. We investigated whether talin-1 and its phosphorylation at Ser 425 play pivotal roles in the activation of integrin heterodimers, which are receptors for ECM components in sEVs.

After talin-1 knockdown, PC3 cells and BxPC3 cells exhibited a significantly diminished ability to spread on ECM components, including fibronectin, laminin, and collagen type I (Fig. 7, A-C). These results suggest that talin-1 in these cells play a pivotal role in mediating cell adhesion via integrins and the ECM. sEVs derived from talin-1 knockdown (KD) or overexpressed (OE) PC3 cells were isolated by ultracentrifugation, and we quantitatively analyzed the numbers of sEV-Halo7-integrinβ1-SF650T and sEV-CD63-Halo7-SF650T particles that bound to ECM components on laminin-coated glass (Fig. 7, D-J). The expression level of talin-1 in sEVs was reduced to 40% following talin-1 KD (top in Fig. 7D). Notably, we observed that the binding of talin-1 KD cell-derived sEVs to laminin was similar to that of sEVs derived from intact cells (top in Fig. 7E; 7F; top in 7H; 7I). Furthermore, overexpression (OE) of talin-1 in sEVs (2.3 times greater than the endogenous level; bottom in Fig. 7D) did not alter the number of sEVs bound to laminin (bottom in Fig. 7E; 7G; bottom in 7H; 7J). Moreover, the laminin-binding affinity of sEVs derived from talin-1 KD BxPC3 cells, in which the talin-1 content was reduced to 11% of that in sEVs from wild-type cells (Fig. 7K), was not significantly different from that of sEVs derived from wild-type cells (Fig. 7, L and M). These results suggest that the amount of talin-1 in sEVs is irrelevant to the binding of sEVs to laminin on glass.

**Fig. 7.**
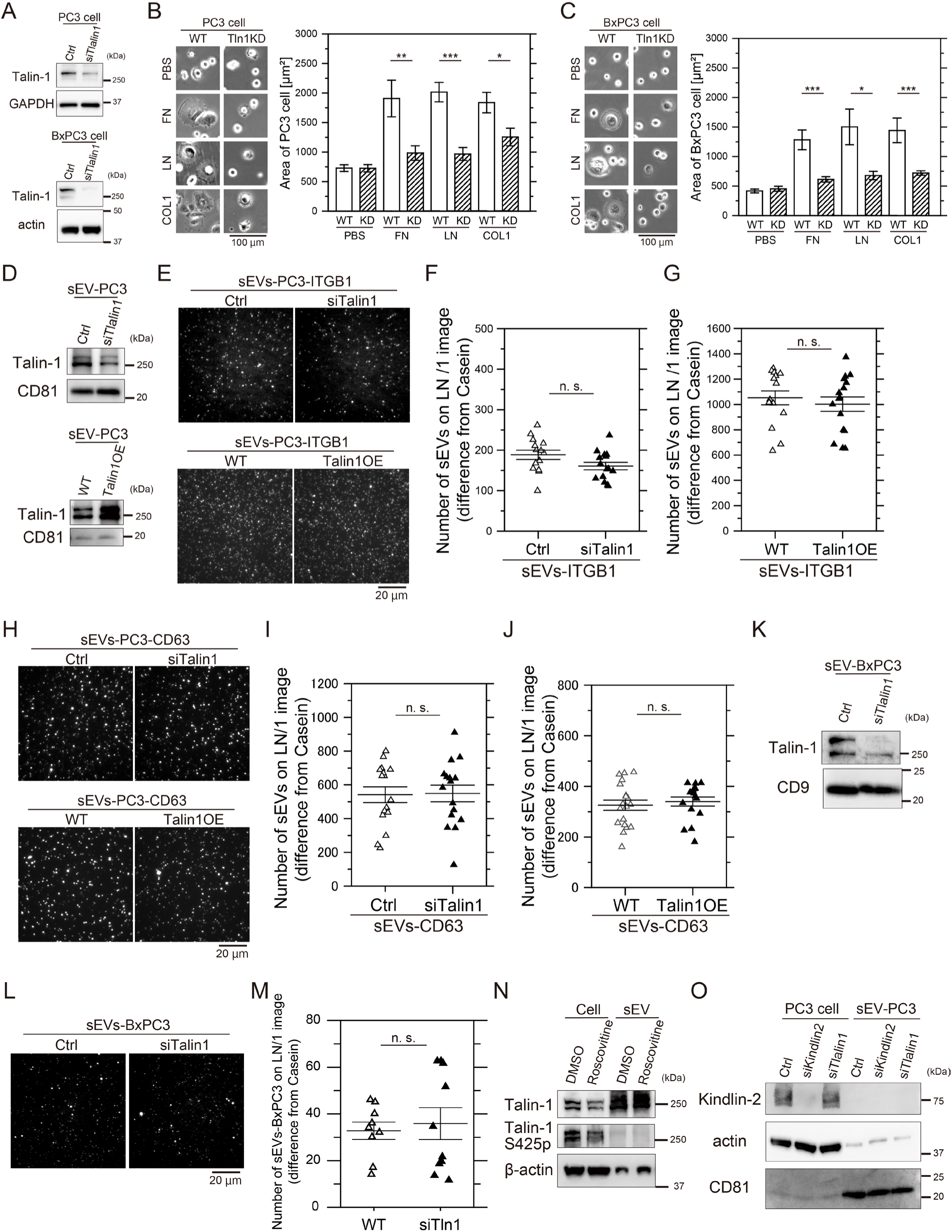
Talin-1 in sEVs does not regulate the binding affinity of integrins for laminin. (A) Western blot analysis of PC3 and BxPC3 cells after talin-1 knockdown (KD) by siRNA. (B, C) Cell spreading assay of wild-type (WT) and talin-1 (Tln1)-KD PC3 cells (B) and BxPC3 cells (C) on glass coated with ECM components: fibronectin (FN), laminin (LN), or collagen typeⅠ (COL1). Cells were observed after 2 hours of incubation, and cell areas were quantified. (D) Western blot analysis of PC3 cell-derived sEVs after talin-1 KD by siRNA or overexpression of talin-1. (E, F, and G) The fluorescence images (E) and the numbers of PC3-sEVs attached to glass coated with laminin before and after (F) talin-1 KD or (G) overexpression of talin-1. Halo7-integrin β1 in sEVs was labeled with SF650T. (H, I, and J) The fluorescence images (H) and the numbers of CD63-labeled sEVs attached to glass coated with laminin before and after (I) talin-1 KD and (J) overexpression of talin-1. (K) Western blot analysis of BxPC3 cell-derived sEVs after talin-1 KD by siRNA. (L and M) Fluorescence images and the numbers of sEVs-BxPC3 bound to laminin on glass before and after talin-1 KD. The membranes of sEVs were stained by Exosparkler DeepRed. (N) Western blot analysis of the phosphorylation of Ser425 on talin-1 in PC3 cells and sEVs. Roscovitine: an inhibitor of CDK5 that phosphorylates Ser425 of talin-1. (O) Western blot analysis of kindlin-2 in PC3 cells and PC3-derived sEVs before and after kindlin-2 KD and talin-1 KD. Data are presented as the mean ± SE. n.s., non-significant difference; \**P*<0.05; \*\**P*<0.01; \*\*\**P*<0.001 according to Welch’s t-test (two-sided).

Subsequently, we investigated whether the phosphorylation of talin-1 at Ser 425 promotes the binding of integrin heterodimers in sEVs to ECM components. Western blotting revealed that talin-1 was highly phosphorylated at Ser 425 in PC3 cells, but this phosphorylation was attenuated by treatment with roscovitine, an inhibitor of Cdk5 (Fig. 7N). Moreover, talin-1 at Ser 425 was almost completely dephosphorylated in sEVs (Fig. 7N), suggesting that the phosphorylation of talin-1 at Ser 425 does not play an important role in the binding of sEVs to ECM components. Furthermore, kindlin-2, a co-activator of integrins abundantly expressed in cancer cells (Montanez et al., 2008), was not detected in sEVs (Fig. 7O). Combined with the finding from talin-1 KD/OE and Ser425 dephosphorylation experiments, these results unequivocally indicate that talin-1 and kindlin-2 in sEVs do not increase the binding of integrin heterodimers in sEVs to laminin. Moreover, the diminished binding of integrin heterodimers to fibronectin in EVs might stem from the absence of talin-1 and kindlin-2 functionality.

### Cholesterol attenuates the binding of sEVs to laminin and the PMs of recipient cells

Given that our results demonstrate that talin-1 and kindlin-2 do not participate in the activation of integrin in sEVs (Fig. 7), we tried to identify alternative factors that specifically maintain integrin binding to laminin even in sEVs where inside-out signaling is lacking. Previous reports have shown that focal adhesions are highly ordered structures similar to rafts (Gaus et al., 2006) and that some integrin subunits, such as α6 and β4, undergo palmitoylation and partition into rafts (Gagnoux-Palacios et al., 2003; Yang et al., 2004). Moreover, a recent study demonstrated that sEV membranes are richer in rafts than the parental cell PM (Yasuda et al., 2022). Therefore, we hypothesized that rafts in sEV membranes enhance the binding of integrins in sEVs to laminin. To test this possibility, we used TIRFM to observe the binding of sEV-CD63-Halo7-SF650T to laminin on glass after depletion or addition of cholesterol. After cholesterol depletion by MβCD or saponin, the number of sEVs that bound to laminin on glass significantly increased by factors of 1.7 and 1.3, respectively (p < 0.01) (Fig. S6, A and B). Conversely, after cholesterol was added to sEVs with the MβCD-cholesterol complex, the binding affinity was significantly reduced by approximately 20% (p < 0.01) (Fig. S6C). Furthermore, we quantified the binding affinity of cholesterol-depleted and cholesterol-supplemented sEVs for MRC-5 PMs. In agreement with the findings on glass, cholesterol depletion from sEVs significantly increased the number of sEVs that bound to the recipient cell PM after 30 min of incubation (p = 0.039), whereas the addition of cholesterol markedly reduced the binding (p = 0.0059) (Fig. S6, D-F). Thus, these results explicitly show that, in contrast to our hypothesis, the presence of abundant cholesterol in sEV membranes impedes the binding of sEVs to both laminin and the recipient cell PM.

### CD151 preserves the binding of integrin heterodimers in sEVs to laminin

Since cholesterol inhibits the activity of laminin receptors in sEVs, we investigated the role of a membrane molecule that interacts with both cholesterol and integrin heterodimers in laminin binding in sEVs lacking inside-out signaling. CD151 is a ubiquitously expressed tetraspanin protein, and its large outer loop (Kazarov et al., 2002) specifically interacts with the C-terminal domain of the extracellular region of the integrin α7, α6, and α3 subunits, selectively strengthening the binding of integrin heterodimers such as α6β1 and α6β4 to laminin (Lammerding et al., 2003; Winterwood et al., 2006). CD151 has a cholesterol-binding domain (Purushothaman and Thiruvenkatam, 2019), is palmitoylated (Yang et al., 2002), and associates with many other membrane proteins through raft-lipid interactions (Berditchevski, 2001; Charrin et al., 2003; Odintsova et al., 2006). Therefore, we investigated whether the binding activities of integrin heterodimers in sEVs lacking inside-out signaling to laminin are preserved by CD151. After CD151 KD by siRNA (Fig. 8A), there was no change in the levels of integrin α6 and α3 in PC3 cells. However, these cells exhibited a significantly reduced ability to spread on laminin, whereas their ability to spread on fibronectin and collagen type I remained unaltered (Fig. 8B). CD151 KD also decreased the number of sEVs that bound to laminin by 41% (Fig. 8C). Furthermore, CD151 KD attenuated the marked increase in the laminin-binding activity of sEVs caused by cholesterol depletion (Fig. 8, C and D). These results suggest that the presence of cholesterol in sEVs suppresses sEV binding to laminin through a CD151-dependent mechanism. In other words, an unidentified molecule that interacts with CD151 in a cholesterol-dependent manner may inhibit the binding of integrin heterodimers in sEVs to laminin.

**Fig. 8.**
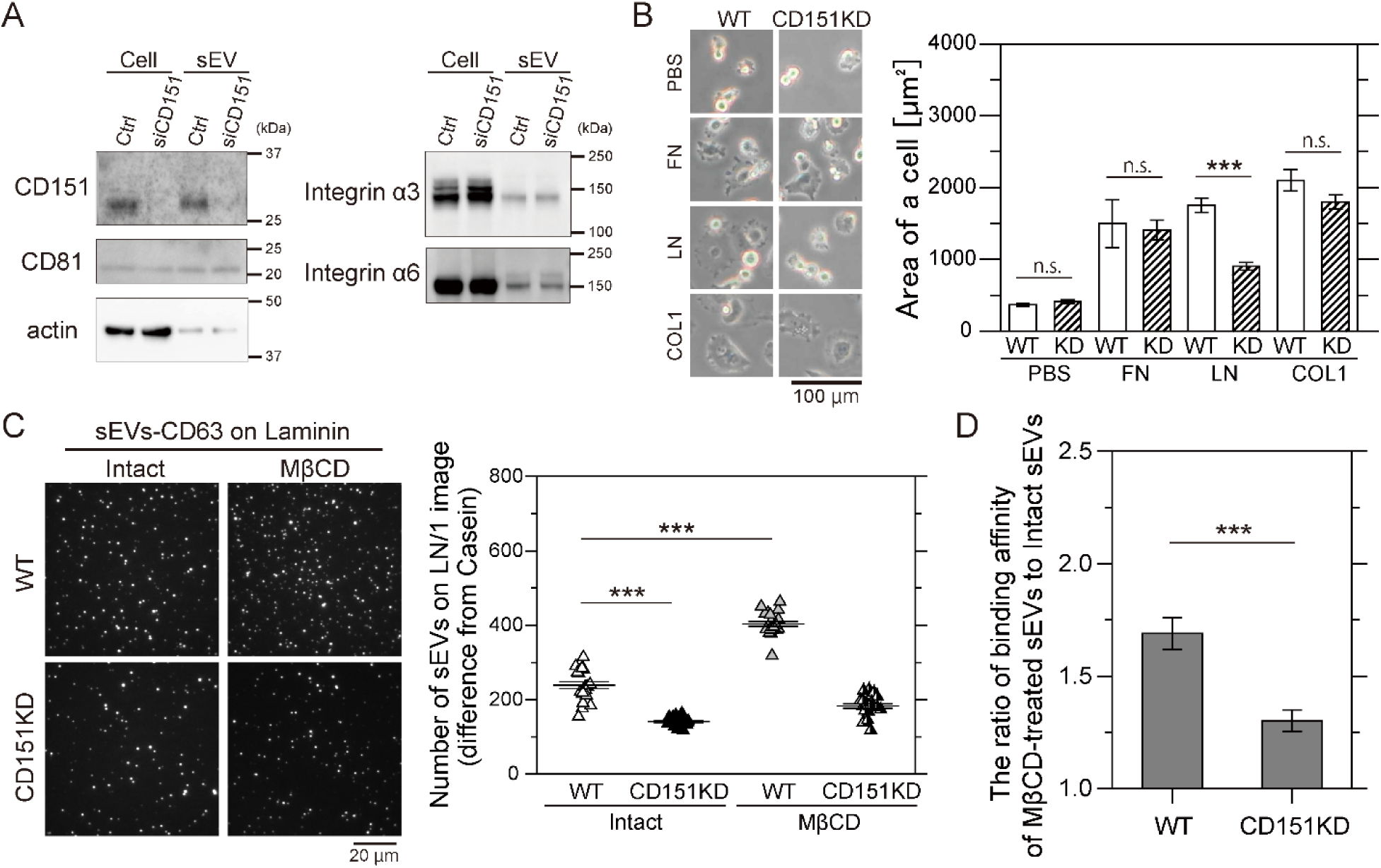
CD151 and cholesterol regulate the binding affinity of sEVs for laminin. (A) Western blot analysis of CD151 and integrin subunits in PC3 cells and sEVs after CD151 KD. The amount of cell proteins loaded in one lane was 2.5 times greater than that of the sEVs in the other lane. (B) Images of wild-type (WT) PC3 cells and CD151-knockdown cells on glass coated with ECM components (fibronectin (FN), laminin (LN), or collagen type Ⅰ (COL1)) after 2 h of incubation. The areas of the cells were quantified. (C) Single-particle fluorescence images of sEV-CD63Halo7-SF650T particles bound to laminin (LN) on glass before and after CD151 was knocked down and cholesterol was depleted by MβCD (left). The number of attached sEVs increased (right). (D) The binding affinity ratio of cholesterol-depleted sEVs to intact sEVs was compared with that of CD151 KD sEVs. Data are presented as the mean ± SE. n.s., non-significant difference; \**P*<0.05; \*\**P*<0.01; \*\*\**P*<0.001 according to Welch’s t-test (two-sided).

### PC3-derived sEVs induce endothelial branching morphogenesis of HUVEC in a laminin-dependent manner

We next examined whether laminin- and integrin-mediated binding of sEVs to recipient cells play a role in cellular responses. Prostate cancer-derived sEVs have been reported to promote an angiogenic phenotype in human umbilical vein endothelial cells (HUVEC) (Progol et al., 2021; Wang et al., 2024). Thus, we investigated whether laminin-mediated binding of PC3-derived sEVs contributes to the morphological branching in HUVEC associated with angiogenesis (Myers et al., 2011). To determine whether laminin is essential for PC3-derived sEV binding to HUVEC, we knocked down laminin γ1 in HUVEC. As laminin γ1 is highly abundant among γ chain isoforms in HUVEC, according to the *Human Protein Atlas* (https://www.proteinatlas.org/), its knockdown should also reduce laminin α subunits (Fleger-Weckman et al., 2016). Indeed, Western blotting analysis confirmed that siRNA-mediated reduction of laminin γ1 to 13% of its original level (top in Fig. 9A) resulted in a corresponding decrease in total laminin to 34% (middle in Fig. 9A). The number of sEVs-PC3-CD63Halo7-TMR particles bound to both the apical and basal membranes of HUVEC, quantified by confocal fluorescence microscopy (Fig. 9B), was dramatically decreased to 41% of the original level after laminin γ1 KD (Fig. 9C). Similarly, the number of sEVs bound to the basal PM, observed by TIRFM, was markedly lower in laminin γ1 KD-HUVEC than wild-type cells at prolonged incubation (> 30 min) (Fig. 9D). To assess morphogenesis, HUVECs were treated with PC3-derived sEVs for 12 hours, and their branched protrusions were examined using TIRFM. sEV-treated HUVEC exhibited significantly elongated protrusions compared to untreated cells (top-left and top-middle panels in Fig. 9E). Quantitative analysis of average protrusion lengths (yellow bars, Fig. 9F) (Myers et al., 2011; Braun et al., 2014) in all examined cells and average total protrusion length per cell revealed that laminin γ1 KD significantly suppressed protrusion elongation (bottom-middle panel in Fig.9E and Fig. 9, G-J), demonstrating that laminin-mediated sEV binding induces these morphological alterations. Interestingly, HUVEC stimulation with 0.26 nM VEGF, an angiogenesis-inducing molecule, for 12 hours produced protrusion lengths comparable to those observed after laminin γ1 KD (top-right and bottom-right in Fig. 9E and Fig. 9, G-J). These results indicate that while laminin is not intrinsically required for HUVEC branching morphogenesis, the binding of sEVs to laminin is essential for sEV-induced morphological changes in HUVEC.

**Fig. 9.**
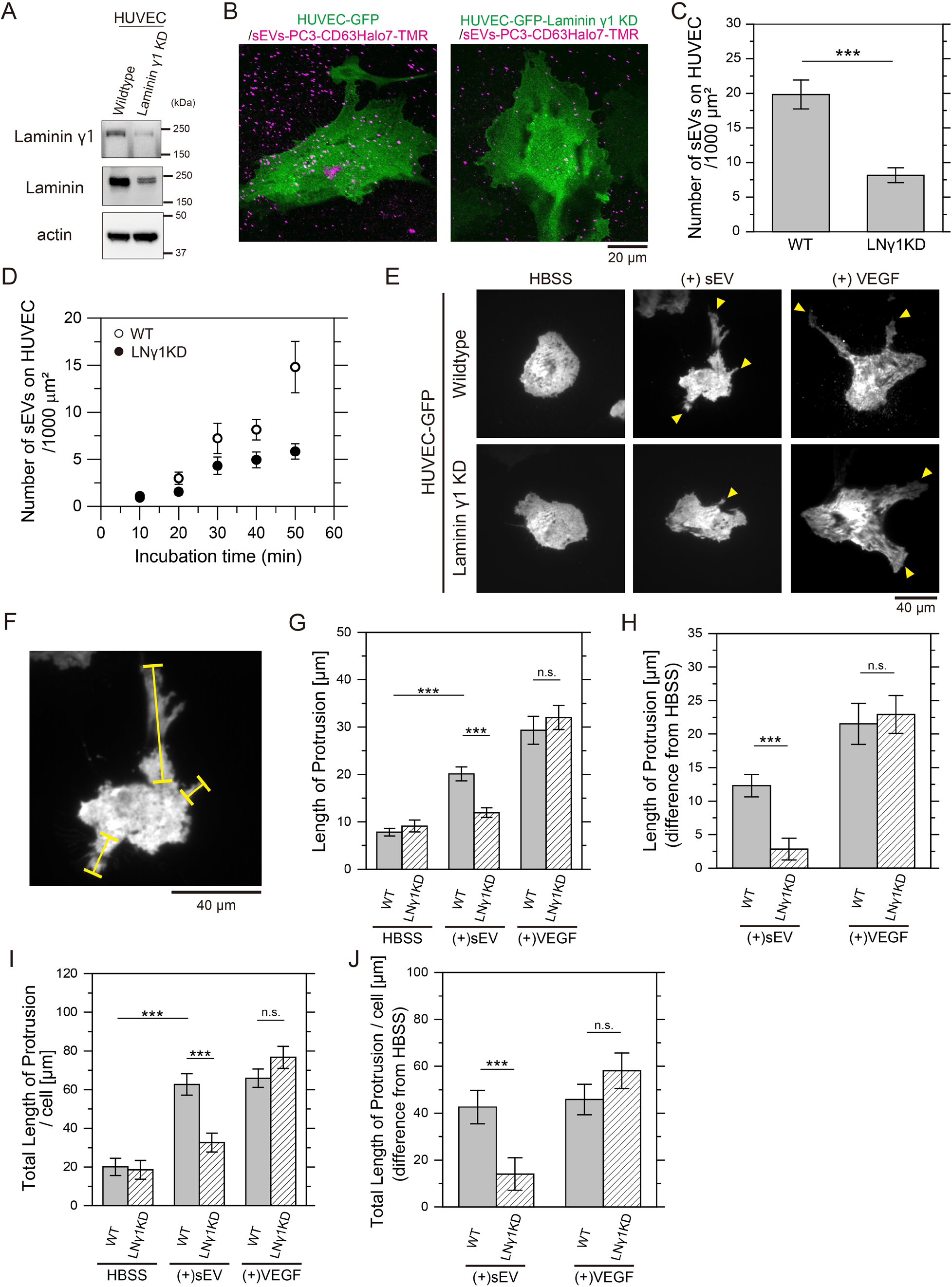
Laminin-mediated binding of PC3-derived sEVs induces endothelial morphogenesis in HUVEC. (A) Western blot analysis of laminin γ1 and total laminin levels in laminin γ1-KD HUVEC. (B) Fluorescence images of wild-type (WT) or laminin γ1-KD HUVECs expressing GFP and sEVs-PC3-CD63Halo7-TMR bound to the cells, observed by confocal microscopy. Cells were fixed after 1-hour incubation with sEVs. (C) Quantification of sEV particles bound to both apical and basal PM of WT or laminin γ1-KD HUVEC by confocal microscopy. (D) Time course analysis of the number of sEV particles bound to the basal PM of live WT or laminin γ1-KD HUVEC, monitored using TIRFM. (E) Fluorescence images of HUVEC expressing GFP after 12 hours of treatment with PC3-derived sEVs or 10 ng/ml VEGF. Yellow arrowheads show branched protrusions of HUVEC. (F) Protrusion lengths measured along yellow lines as indicated. (G) Average protrusion lengths across all examined cells before and after treatment with sEVs or VEGF. (H) Changes in average protrusion length after treatment with sEVs or VEGF, relative to untreated conditions. (I) Average total protrusion length per cell before and after treatment with sEVs or VEGF. (J) Variations in average total protrusion length per cell after treatment with sEVs or VEGF, relative to untreated conditions. Data are presented as mean ± SE. n.s., non-significant difference; \**P*<0.05; \*\**P*<0.01; \*\*\**P*<0.001 according to Welch’s t-test (two-sided).

## Discussion

Our results demonstrated that laminin, not fibronectin, is the primary target of tumor-derived sEV subtypes containing CD63, CD81, or CD9. The binding of sEVs to laminin is mediated by integrin α6β1 and α6β4 heterodimers, and GM1 in sEVs, as evidenced by experiments performed on glass (Fig. 2, Fig. S2 and Fig. S4) and recipient cell PMs (Fig. 4, Fig. 6 and Fig. 10A). Tumor cell-derived mEVs and MVs also displayed higher binding affinities for laminin than fibronectin (Fig. 3, C-E), despite containing receptors for both (Fig. 1A and Fig. 3F). Therefore, factors beyond ECM receptor presence likely influence binding activities. While talin-1 and kindlin-2 do not enhance integrin binding activity in EVs (Fig. 7), the tetraspanin CD151, considerably increased integrin-laminin binding, whereas cholesterol suppressed receptor function in a CD151-dependent manner. (Fig. 8). The binding of tumor-derived sEVs induced endothelial branching morphogenesis in HUVEC in a laminin-dependent manner (Fig. 9).

**Fig. 10.**
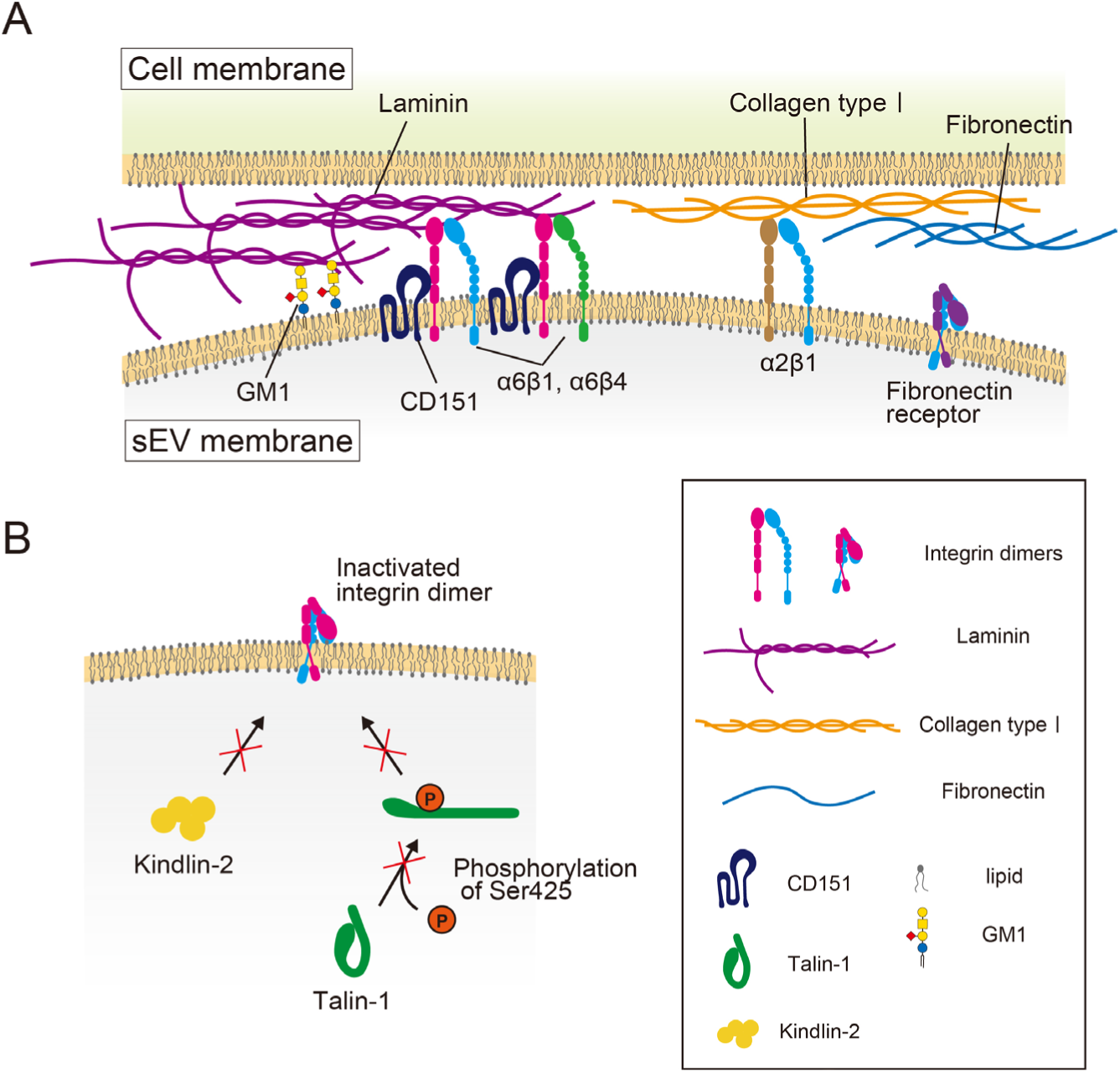
Schematic model showing the binding of EVs to the recipient cell PM. (A) Integrin heterodimers in EVs, α6β1/α6β4 and α2β1, can bind to laminin and collagen type I, respectively, but integrin heterodimers in EVs, such as α5β1, bind only weakly to fibronectin. CD151 enhances the binding of integrins α6β1 and α6β4 in sEVs to laminin. Cholesterol in sEVs suppresses laminin binding through a CD151-dependent mechanism. (B) Because Ser 425 of talin-1 is hardly phosphorylated in EVs, talin-1 is inactive and is not involved in promoting the ECM-binding activity of integrin heterodimers. Kindlin-2 is also not involved in the activation of integrin heterodimers in EVs.

Although the inhibition of EV binding to recipient cells by antibodies and ligands against integrins has suggested that integrin subunits in EVs are responsible for the binding of EVs to recipient cells (Nazarenko et al., 2010; Carney et al., 2017; Zhang et al., 2022), there has been no direct evidence of the specific binding of integrins to ECM components. The binding affinities of EVs for fibronectin, laminin, and collagen type I have not been quantitatively compared. In this study, we established an in vitro experimental system to quantitatively measure the binding affinities of integrin heterodimers with ECM components, which enabled us to compare the binding affinities of EVs derived from different types of tumor cells with fibronectin, laminin, and collagen type I. The strong binding of all the sEVs, mEVs and MVs to laminin via integrin heterodimers and the weak binding to fibronectin were unexpected because previous studies reported that the binding of EVs to recipient cells was inhibited by antibodies and inhibitors against fibronectin receptors (Huang et al., 2016; Altei et al., 2020). However, since our results did not indicate that tumor-derived EVs have no binding ability to fibronectin at all but rather that they exhibit lower binding to fibronectin than to laminin, our results are not inconsistent with previously reported findings. Since the amount of EVs bound to PMs increases with the amount of EVs added to cells, differences in the numbers of EVs bound to PMs after treatment with antibodies against integrin subunits of fibronectin receptors may be detected in previous reports.

Our results also revealed that only GM1, which is approximately 29-fold more enriched in sEVs compared to sEV-secreting cells, bound to laminin on glass, while all the other examined gangliosides did not (Fig. S4). Furthermore, a competitive binding inhibition assay using the GM1 glycan moiety revealed that 18% and 61% of sEVs were bound to laminin through GM1-laminin and integrin β1-laminin interactions, respectively (Fig. S4L). Meanwhile, none of the examined gangliosides, including GM1, bound to fibronectin. These results suggest that gangliosides may bind to laminin through specific glycan–protein interactions, not merely through electrostatic interactions. Since the binding of the glycan structure of GM1 to laminin does not require complicated regulation by other membrane molecules, it must constitute a robust binding system that is not prone to loss of activity unless GM1 is degraded.

Integrin heterodimers in cell membranes transition between bent-closed and extended-open conformations, maintaining the extended-open state by binding to ligands or activators like talin (Moser et al., 2009; Li et al., 2024). Our study demonstrates that integrin β1 in sEVs adopts the extended-open conformation when bound to laminin (Fig. 5C). Structural analysis revealed that integrin α6β1 binds laminin γ1’s C-terminal tail via the β-propeller domain of integrin α6, with R155 being crucial for this interaction (Arimori et al., 2021). Mutations at R155 in integrin α6 significantly reduced sEV binding to laminin (Fig. 5F), indicating the reliance on conventional integrin-ligand interactions. These findings suggest that integrin heterodimers in sEVs, like those in cells, bind laminin through the extended-open conformation and that sEV integrin function may be modulated by established integrin-regulating molecules.

Neither the knockdown nor the overexpression of talin-1 in sEVs significantly altered the binding of sEVs to laminin (Fig. 7). In addition, Ser 425 of talin-1, an activating phosphorylation site (Jin et al., 2015), was almost completely dephosphorylated, whereas Ser 425 was phosphorylated in cells that secreted sEVs. Talin-1 is ubiquitously expressed in all cell types in all tissues, whereas talin-2 is expressed mostly in the brain, heart muscle, and kidney (Monkley et al., 2001; Thul et al., 2017) and is not expressed in the PC3 cells we examined. Moreover, kindlin-2 was not detected in sEVs (Fig. 7O). These results explicitly demonstrate that inside-out signaling by integrins is absent in sEVs and that integrins in sEVs exist in a completely different environment from that in cell PMs. The lack of inside-out integrin signaling in sEVs seems reasonable because there is no cytoskeletal actin filament network inside the sEVs, which might be one of the reasons that integrin heterodimers of fibronectin receptors exhibit relatively low activity in sEVs (Fig. 10B). On the other hand, our results indicate that α6β1 and α6β4 integrin heterodimers of laminin receptors in sEVs are still active (Fig. 2, Fig. 4, Fig. 6, and Fig. S2), and a tetraspanin, CD151, preserves the activities of these receptors (Fig. 8C). CD151 is known to form a complex with integrin early in biosynthesis; this complex is delivered to tetraspanin-enriched microdomains (TEMs) in the cell PM (Stipp, 2010) and selectively strengthens integrin-mediated adhesion to laminin (Lammerding et al., 2003). CD151 has a cholesterol-binding domain (Purushothaman and Thiruvenkatam, 2019) and links integrin heterodimers of laminin receptors with other tetraspanins by cholesterol- and raftophilic lipid-dependent mechanisms (Berditchevski et al., 2002; Franco et al., 2010). Our results revealed that cholesterol inhibited the binding of integrin heterodimers in sEVs to laminin in a CD151-dependent manner (Fig. 8, C and D), suggesting that the CD151-dependent activation of integrin heterodimers in sEVs may be attenuated by the association of CD151 with lipid rafts. This finding is consistent with the fact that the depalmitoylation of CD151 increases the downstream signaling of integrins in the cell PM (Berditchevski et al., 2002). Since the PC3-derived sEV membrane possesses a greater abundance of the Lo (raft)-like phase than the cell PM (Yasuda et al., 2022), attenuation of CD151-dependent activation of integrin heterodimers may occur more extensively in sEVs than in cell PMs.

The higher binding affinity of tumor-derived sEVs for laminin over fibronectin on recipient cell PMs may influence recipient cell response. For example, embryonic neural crest cells (NCCs) exhibit ligand- and receptor-specific integrin regulation (Strachan and Condic, 2004). On laminin-coated glass, NCCs accumulate internalized laminin receptors (integrin α6) but not fibronectin receptors (integrin α5). Internalized laminin receptors colocalize with recycling vesicles and return to the cell surface, whereas fibronectin receptors do not. Blocking receptor recycling inhibited NCC motility on laminin, indicating that laminin receptor recycling is essential for rapid migration. Recent PALM movie imaging and single-particle tracking revealed that sEVs bound to laminin on recipient cells accumulate integrin subunits underneath, which are internalized via caveolae, and clathrin- and caveolae-independent pathways. Approximately 20-40% of PC3 cell-derived sEVs accumulate in recycling endosomes (Hirosawa et al., 2024, *Preprint*), suggesting that sEV-laminin binding promotes efficient integrin recycling.

Another example of a specific response to integrin-laminin binding involves signaling pathways distinct from those induced by fibronectin. Laminin-10/11 (same as laminin 511 used in this study) activates Rac via p130Cas-CrkII-DOCK180, enhancing cell migration, while fibronectin activates Rho via FAK, reducing cell migration (Gu et al., 2001). Since sEV binding to recipient cell PMs rapidly activates integrin, talin-1, and Src family kinases (Hirosawa et al., 2024, *Preprint*), our results suggest laminin-specific signaling may occur during sEV binding.

Intriguingly, our results revealed that the binding of PC3-derived sEVs to HUVEC induced branching morphogenesis in a laminin-dependent manner, comparable to the level induced by VEGF in a laminin-independent manner (Fig. 9). As these morphological changes occurred 12 hours after sEV treatment, they are likely driven by internalized sEVs rather than prompt sEV-induced signaling at the PM. A previous study suggested that PC3-derived EVs promote branching morphogenesis in HUVEC, potentially mediated by microRNA miR-27a-3p (Prigol et al., 2021). For these cellular responses to occur, the microRNA must first be internalized into HUVEC and subsequently released from internalized EVs into the cytosol. Given that laminin knockdown significantly decreased the binding of sEVs to HUVEC, the amount of internalized microRNA available for cytosolic release is also likely diminished. Another critical factor contributing to the attenuated morphological changes after laminin knockdown may involve alterations in subcellular transport pathways post-internalization, as previously discussed, potentially impairing the efficiency of microRNA release into the cytosol. The mechanisms underlying the enhanced cellular responses induced by laminin-bound EV internalization warrant further investigation.

In summary, our findings demonstrate that sEVs bind to laminin through robust mechanisms that do not require inside-out integrin signaling, leading to laminin-dependent morphological changes in HUVEC. The mechanism of EV binding to laminin revealed in this study will also provide useful information for applied research, including clinical applications.

## Materials and methods

### Materials

The human prostate cancer cell line (PC3), human breast cancer cell line (SKBR3), human pancreas cancer cell line (BxPC-3), human fetal lung fibroblast line (MRC-5), and human marrow stromal cell line (HS-5) were purchased from the American Type Culture Collection (ATCC). The human breast cancer cell line (4175-LuT) was kindly provided by Dr. Joan Massagué and Dr. Hoshino (Hoshino et al., 2015). HUVEC (human umbilical vein endothelial cells) were purchased from PromoCell.

All antibodies were obtained from commercial sources. The following primary antibodies were used for western blotting: mouse monoclonal anti-beta actin 15G5A11/E2 (1:5000, Thermo Fisher, Cat# MA1-140), mouse monoclonal anti-CD63 8A12 (1:1000, Cosmo Bio, Cat# SHI-EXO-M02), mouse monoclonal anti-CD81 B11 (1:500, Santa Cruz Biotechnology, Cat# SC166029), rabbit monoclonal anti-CD9 EPR23105-125 (1:1000, Abcam, Cat# ab263019), mouse monoclonal anti-CD151 11G5a (1:250, Abcam, Cat# ab33315), rabbit polyclonal anti-fibronectin (1:1000, Sigma‒Aldrich, Cat# F3648), rabbit monoclonal anti-GAPDH 14C10 (1:2000, Cell Signaling, Cat# 2181S), rabbit monoclonal anti-integrin alpha 2 EPR5788 (1:2500, Abcam, Cat# ab133557), rabbit polyclonal anti-integrin alpha 3 (1:500, Abcam, Cat# ab190731), rabbit polyclonal anti-integrin alpha 5 (1:250, Cell Signaling, Cat# 4705S), rabbit monoclonal anti-integrin alpha 5 clone EPR7854 (1:500, Abcam, Cat#ab150361), rabbit polyclonal anti-integrin alpha 6 (1:500, Cell Signaling, Cat# 3750S), rabbit polyclonal anti-integrin alpha 7 (1:500, Abcam, Cat# ab203254), rabbit polyclonal anti-integrin alpha V (1:500, Abcam, Cat# ab117611), mouse monoclonal anti-CD29 Clone 18 (1:2000, BD Bioscience, Cat# 610467), rabbit polyclonal anti-integrin beta 3 (1:500, Millipore, Cat# AB2984), rabbit polyclonal anti-integrin beta 4 (1:500, Cell Signaling, Cat# 4707S), rabbit polyclonal anti-integrin beta 5 (1:500, Cell Signaling, Cat# 4708S), rabbit polyclonal anti-laminin 1+2 (1:1000, Abcam, Cat# ab7463), mouse monoclonal anti-laminin clone 2E8 (1:500, Millipore, Cat#MAB1920), mouse monoclonal anti-talin-1 97H6 (1:500, Gene Tex, Cat# GTX38972), rabbit polyclonal anti-phospho-talin (Ser-425), (1:250, ECM Biosciences, Cat# TP4171), mouse monoclonal anti-kindlin-2 clone 3A3 (1:500, Sigma-Aldrich, Cat# MAB2617), and anti-GM3 antibody clone GMR6 (1:1000, Tokyo Chemical Industry, Cat# A2582), along with biotin-conjugated Cholera toxin B subunit (1:500, List Labs, Cat# 112). Additionally, goat anti-mouse IgG-HRP (1:5000, Millipore, Cat# 12-349), donkey anti-rabbit IgG-HRP (1:4000, Cytiva, Cat# NA934), goat anti-rabbit IgG-HRP (1:10000, Sigma‒Aldrich, Cat# A0545), goat anti-mouse IgM antibody-HRP (1:10000, Invitrogen, Cat# 31440), and rabbit anti-biotin antibody-HRP clone D5A7 (1:10000, Cell Signaling, Cat# 5571) were used as secondary antibodies. For immunoprecipitation, mouse monoclonal IgG2a negative control C1.18.4 (Millipore, Cat# MABF1080Z), mouse monoclonal IgG1-κ negative control MOPC-21 (Millipore, Cat# MABF1081Z), and rabbit polyclonal IgG negative control (Abcam, Cat# ab37415) were used. For the observation of EVs and ECM components, mouse monoclonal anti-CD81 JS-81 (BD Pharmingen, Cat# 555675), mouse monoclonal anti-CD9 M-L13 (BD Pharmingen, Cat# 555370), rabbit polyclonal anti-collagen type 1 (Abcam, Cat# ab34710), rabbit polyclonal anti-fibronectin (Abcam, Cat# ab23750), rabbit polyclonal anti-laminin (Sigma‒Aldrich, Cat# L9393), goat polyclonal anti-rabbit IgG-rhodamine (Cappel, Cat# 55666), and goat anti-rabbit IgG (Cappel, Cat# 0212-0081) were used.

We purchased the following purified ECM proteins: fibronectin from Roche (Cat# 11051407001), laminin from Sigma‒Aldrich (Cat# L6274), and collagen type I from Sigma‒ Aldrich (Cat# C7774). For the experiments involving cholesterol and CD151, we used laminin-511 (Biolaminin 511 LN, BioLamina). The recombinant laminin (L6274, Sigma‒Aldrich) we used was almost identical to Biolaminin 511 LN because nearly all the laminins in L6274 were laminin-511 (Wondimu et al., 2006).

### Cell culture

Mycoplasma contamination was not detected in any of the cell lines used in this study.

PC3 human prostate cancer cells (ATCC, CRL-1435) were cultured in HAM F12 medium (Sigma‒Aldrich) supplemented with 10% fetal bovine serum (FBS) (Gibco), 100 U/ml penicillin, and 100 μg/ml streptomycin (Gibco). Human fetal lung fibroblasts (MRC-5; ATCC, CCL-171) and human uterine cervix cancer cells (HeLa) were cultured in minimum essential medium (Gibco) supplemented with 10% FBS, 100 U/ml penicillin, and 100 μg/ml streptomycin. Human marrow stromal cells (HS-5; ATCC, CRL-11882) and human breast cancer cells (4175-LuT; Hoshino et al., 2015) were cultured in Dulbecco’s modified Eagle’s medium (Sigma‒ Aldrich) supplemented with 10% FBS, 100 U/ml penicillin, and 100 μg/ml streptomycin. Human breast cancer cells (SKBR3; ATCC, HTB-30) were cultured in McCoy’s 5a medium (Gibco) supplemented with 10% FBS, 100 U/ml penicillin, and 100 μg/ml streptomycin. Human pancreatic cancer cells (BxPC-3; ATCC, CRL-1687) were cultured in RPMI-1640 medium (Sigma‒Aldrich) supplemented with 10% FBS, 100 U/ml penicillin, and 100 μg/ml streptomycin. HUVEC were cultured in Endothelial Cell Growth medium (PromoCell).

The mouse melanoma cell line B78 was cultured in Dulbecco’s modified Eagle’s medium (Sigma‒Aldrich) supplemented with 10% FBS, 100 U/ml penicillin, 100 μg/ml streptomycin and 400 μg/ml G418. B78 cells expressed only GM3 (Fig. S4). pM2T1-1 cDNA transfection induced the overexpression of GM2/GD2 synthase, resulting in GM3 depletion because this enzyme uses GM3 as a substrate for synthesizing GM2 (Fig. S4). The transfection of B78 cells with pM1T-9 and pM2T1-1 induced the overexpression of GM1 (Fig. S4), resulting in GM3 and GM2 depletion because these enzymes use GM3 and GM2 as substrates for the synthesis of GM1. Transfection of B78 cells with pM2T1-1 and subsequently with pD3T31 induced overexpression of GD2, while transfection with pD3T31 and subsequently with pM2T1-1 induced overexpression of both GD2 and GD3 (Fig. S4) (Yesmin et al., 2023).

### Flow cytometry of cells expressing gangliosides

The expression of gangliosides on the cell surface was analyzed by flow cytometry with a FACSCalibur (Becton-Dickison) as previously reported (Yamashiro et al., 1995; Yesmin et al., 2023). Intact and glycosyltransferase-transfected B78 cells were stained by incubation with monoclonal antibodies against GM3 (M2590, IgM) [Cosmo Bio], GM2 (10-11, IgM), GD2 (220-51, IgG1), GD3 (R24, IgG3) [10-11 and R24 antibodies kindly provided by P. O. Livingston and L. J. Old at Memorial Sloan-Kettering Cancer Center], GD1a (D-266, IgM), or GD1b (370, IgM); FITC-labeled goat anti-mouse IgM antibodies (Cappel); or goat anti-mouse IgG (H & L chain) antibodies (Cappel). Anti-GD2, anti-GD1a, and anti-GD1b antibodies were generated by K. Furukawa. Alternatively, GM1 on the cells was stained by applying biotinylated cholera toxin subunit B (CTXB; Sigma‒Aldrich) and then FITC-labeled avidin (Zymed).

### Knockout of integrin subunits

The integrin β1, β4, α2, and α6 subunits were knocked out in PC3 cells via CRISPR–Cas9 gene editing. The single guide RNA (sgRNA) target sequence was selected via the online tool CHOPCHOP (https://chopchop.cbu.uib.no/). The sgRNA target sequences were subsequently incorporated into the multicloning site of pSpCas(BB)-2A-Puro (addgene) using BbsI-HF (New England Biolabs). The sgRNA target sequences used were TCATCACATCGTGCAGAAGT and ATACAAGCAGGGCCAAATTG for the knockout of integrin β1; GTGCAGGTTCTCGTATCCCT and GCGCTCAGTCAAGGTAAGCG for the knockout of integrin α2; CCGGGTACTTATAACTGGAA and ACACCGCCCAAAGATGTCTC for the knockout of integrin α6; and CTGCGAGATCAACTACTCGG for the knockout of integrin β4. PC3 cells were transfected with plasmids with a 4D-Nucleofector (LONZA). The transfected cells were selected and cloned using 1 μg/ml puromycin (final concentration) to achieve the knockout of integrin β1 or α2. For the knockout of integrin α6 or β4, the transfected cells were sorted by flow cytometry (BD, FACSMelody) using a PE-conjugated anti-integrin α6 antibody (Invitrogen) or a PE-conjugated anti-integrin β4 antibody (BD). The knockout of each integrin subunit was validated by western blotting.

### Preparation of cDNA plasmids and transfection

Wild-type PC3 cells and integrin KO PC3 cells were transfected with 1500 ng of cDNA encoding CD63-Halo7, CD81-Halo7, or CD9-Halo7 using a 4D-nucleofector (LONZA) according to the manufacturer’s recommendations. cDNA plasmids for human CD63 (NM_001780), human CD81 (NM_004356 and pFN21AE5228) and human CD9 (NM_001769 and pFN21AE1546) were purchased from Kazusa DNA Res. Inst. The DNA sequences encoding CD63, CD81, or CD9 were cloned and inserted into the pEGFP-N1 vector, and the EGFP sequence was replaced with the Halo7 sequence (Promega). The linker sequence 5’-ACCGGTGGTGGGCGCGCCTCTGGTGGCGGATCCGGGGGT-3’ was incorporated between CD63, CD81, CD9 and Halo7. Cells stably expressing the molecules of interest were selected using a flow cytometer (BD, FACSMelody) and were cloned by treatment with 0.2 mg/ml (final concentration) G418 (Nacalai Tesque). To generate an expression vector for Halo7-integrin β1, cDNA encoding Halo7-integrin β1 was cloned and inserted into the PiggyBac transposon vector. Halo7 was inserted into the hybrid domain of human integrin β1 (NM_002211) between residues Gly101 and Tyr102. The tagging of integrin β1 with Halo7 at this position is known to have no detrimental effects on integrin activity (Huet-Calderwood et al., 2017). Wild-type or integrin β1 KO PC3 cells were cotransfected with cDNA encoding Halo7 integrin β1 and transposase, yielding cell lines that exhibited stable expression of Halo7 integrin β1 (Yusa et al., 2009; Nagai et al., 2021). Cells exhibiting high expression of Halo7-integrin β1 were selected by flow cytometry. cDNA plasmids encoding integrin α6 and the dominant-negative integrin α6 R155A mutant were cloned and inserted into the PiggyBac transposon vector (Yusa et al., 2009). The cDNA plasmid for human integrin α6 (NM_000210.2 and pRK alpha6) was purchased from addgene. The integrin α6 R155A cDNA was generated from the integrin α6 cDNA via PCR. Integrin α6 KO PC3 cells were transfected with cDNA encoding either integrin α6 or the α6 R155A mutant. Stable cell lines expressing these constructs were selected using 0.4 mg/ml zeocin and subsequently cloned.

4175-LuT, PC3, BxPC3, HeLa, and SKBR3 cells were transiently transfected with a plasmid encoding CD63-Halo7 using polyethylenimine (PEI) MAX (Polysciences) to isolate tumor-derived sEVs containing CD63.

To obtain PC3 cells overexpressing human talin-1, PC3 cells were transfected with cDNA encoding human talin-1 (NM_006289) using Lipofectamine 3000 reagent (Invitrogen) according to the manufacturer’s recommendation. cDNA encoding talin-1 was cloned and inserted into the pEGFP-N1 vector, after which the EGFP sequence was deleted. Moreover, talin-1 was knocked down by transfecting PC3 cells and BxPC3 cells with talin-1 siRNA (Santa Cruz Biotech. sc-36610). siRNA-A (Santa Cruz Biotech. sc-37007) was used as a negative control. CD151 or kindlin-2 was knocked down by transfecting PC3 cells with CD151 siRNA (Dharmacon, D-003637-04 and D-003637-13) or kindlin-2 siRNA (Santa Cruz Biotech. Sc-106786) using Lipofectamine 3000.

### Isolation of small extracellular vesicles (sEVs) from cell culture supernatant

Cells that secreted sEVs were cultured either in two 100-mm dishes for observing sEVs on glass or in two 150-mm dishes for western blotting analysis and the observation of sEVs on cell PMs. The cells were grown to approximately 80% confluence (0.3-1.6×10^8^ cells/150-mm dish). These cells were then transfected with plasmids or siRNAs and cultured for 1-2 days. Next, the cell culture medium in the 100-mm or 150-mm dishes was replaced with 10 ml or 30 ml of FBS-free medium, respectively. After 48 h of incubation, the cell culture supernatant was collected. For roscovitine treatment, cells were incubated with 20 μM roscovitine (LKT Laboratories) in FBS-free medium. The collected supernatant was then subjected to centrifugation at 300×g for 10 min at 4°C to remove cells. Subsequently, the supernatant was further centrifuged at 2,000×g and 4°C for 10 min to remove sediment and eliminate apoptotic bodies. Then, the supernatant was centrifuged at 10,000×g and 4°C for 30 min (himac CF16RN, T9A31 angle rotor, 50 ml Falcon tube) to pellet the microvesicles (MVs). Finally, the supernatant was concentrated by ultrafiltration using an Amicon Ultra 15 100K (Millipore) or a Centricon Plus 70 100K (Millipore) filter unit. The collected sEVs were incubated with 50-100 nM (final concentration) HaloTag TMR ligand (Promega) or HaloTag SaraFluor650T (SF650T) ligand (Goryo Chemical) for 1 h at 37°C. The sEVs were then pelleted by ultracentrifugation at 200,000×g and 4°C for 4 h (himac CS100FNX with an S55A2 angle rotor and S308892A microtubes for ultracentrifugation). Alternatively, medium-sized EVs (mEVs) were pelleted by ultracentrifugation at 50,000×g and 4°C for 30 min. The resulting pellet was resuspended in Hank’s balanced salt solution (HBSS) for microscopic observation. For western blotting, the pellet was suspended in RIPA buffer containing protease inhibitor cocktail set III (Millipore).

The ExoSparkler Exosome Membrane Labeling Kit-Deep Red (Dojindo) was also utilized as a fluorescent probe. Microvesicles (MVs) purified by lower-speed centrifugation (10,000×g for 30 min) were successfully labeled according to the Exosparkler Technical Manual. Subsequently, the labeled MVs were purified by ultrafiltration using an Amicon Ultra 0.5 100K filter unit (Millipore).

### Partial cholesterol depletion/addition in sEVs

sEVs were isolated from the cell culture supernatant of two 150-mm dishes using ultracentrifugation. To deplete cholesterol, the sEVs were incubated with 5 mM methyl-beta-cyclodextrin (MβCD) (Sigma‒Aldrich) overnight at room temperature with gentle shaking or incubated with 200 μM saponin (Kanto Chemical) for 1 h on ice. These solutions were then concentrated by ultrafiltration using Amicon Ultra 2 ml 100K. The concentrated sEVs were incubated with 50-100 nM (final concentration) HaloTag SF650T ligand for 1 h at 37°C. Next, the sEVs were pelleted by ultracentrifugation at 200,000×g and 4°C for 4 h. The pellet was resuspended in phosphate-buffered saline (PBS) or RIPA buffer. To add cholesterol, sEVs were incubated with 3.5 mM MβCD-cholesterol complex (1:1) for 2 h at 37°C, then isolated by the same method used in cholesterol depletion. The amount of cholesterol in these sEVs was measured by a LabAssay Cholesterol Kit (Fujifilm Wako Pure Chemical).

### Immunoprecipitation of sEVs

The initial step involved treating 0.6 mg of Protein G-conjugated Dynabeads (Invitrogen) with 200 μl of 15 μg/ml anti-CD63 antibody (Cosmo Bio, SHI-EXO-M02) or mouse IgG2a isotype control (Millipore, MABF1080Z) in PBS containing 0.02% Tween 20. Alternatively, 0.6 mg of Protein A-conjugated Dynabeads (Invitrogen) was treated with 200 μl of 15 μg/ml anti-fibronectin antibody (Sigma‒Aldrich, F3648) or rabbit IgG isotype control (Abcam, ab172730) in PBS containing 0.02% Tween 20. The mixture was incubated with rotation for 10 min at room temperature, and then the supernatant was removed. The beads were washed once with PBS containing 0.02% Tween 20 and twice with 20 mM sodium phosphate (pH 7.0) and 150 mM NaCl using a magnetic rack. Then, the antibody-conjugated beads were resuspended in 5 mM BS3, 20 mM sodium phosphate (pH 7.0), and 150 mM NaCl for crosslinking between the antibody and the beads. After incubation with rotation for 10 min at room temperature, 12.5 μl of 1 M Tris-HCl (pH 7.5) was added to the solution, which was then incubated with rotation for an additional 15 min at room temperature. Subsequently, the supernatant was removed, and the beads were washed once with PBS containing 0.02% Tween 20 and then twice with PBS using a magnetic rack.

The antibody-conjugated beads were mixed with 300 μl of PC3-derived sEVs (0.1 mg/ml protein concentration) in PBS. After the mixture was incubated with rotation overnight, the supernatant was removed, and the beads were washed three times with PBS. Then, the beads were resuspended in Laemmle’s SDS sample buffer and incubated at 95°C for 5 min. The supernatant was collected using a magnetic rack and analyzed by western blotting.

### Transmission electron microscopy of sEVs after negative staining

A copper grid with a carbon-coated acetylcellulose film (EM Japan) was washed briefly (∼1 min). Ten microliters of water was placed on the grid and then blotted to eliminate the excess liquid. Next, 5 μl of the sEV sample was pipetted onto the grid and incubated for 1 min. The excess liquid was then removed by blotting. The grid was then briefly (∼45 sec) treated with 5 μl of 2% phosphor tungstic acid (TAAB Laboratory and Microscopy), followed by blotting to remove excess liquid. Next, the grid was air-dried at room temperature for 3 days in a desiccator containing silica gel desiccant and subsequently subjected to TEM observation.

sEV images were obtained using a single TEM instrument (JEM-2100F, JEOL) at 200 kV. These images were produced by computing the mean of 3-second acquisitions captured by a side-mounted CCD camera (Gatan) and processed by image solution software (Gatan Digital Micrography).

### Determination of sEV size by qNano

The diameter of the suspended sEVs in HBSS was analyzed using a tunable resistive pulse sensing instrument, qNano (Izon Science) (Coumans et al., 2014), according to the manufacturer’s protocol. Izon Control Suite 3.3 (Izon Science) was used for the data analysis. The measurements were performed with an NP100 pore (particle detection range: 40-320 nm) equipped with a stretch of 47.00 mm, a voltage of 1.2 V, and a pressure of 0.8 kPa; an NP150 pore with a stretch of 46.85 mm, a voltage of 1.2 V, and a pressure of 1.4 kPa; or an NP200 with a stretch of 48.00 mm, a voltage of 1.4 V, and a pressure of 1.4 kPa. The samples were calibrated using 70, 95, and 210 nm polystyrene calibration beads (Izom Science, CPC100).

### Western blotting analysis

The quantities of integrin subunits were evaluated in wild-type and integrin-KO PC3 cells, as well as in sEVs derived from these cells, by western blotting. The cells and sEVs were suspended in RIPA buffer containing protease inhibitor cocktail set III (Millipore). The protein concentrations of the samples were determined using a Thermo BCA protein assay kit (Thermo Fisher Scientific) and adjusted to 1 mg/ml for SDS‒PAGE. Then, 5× concentrated Laemmli’s SDS sample buffer was added to the samples, and the mixture was incubated at 95°C for 5 min in a blocking incubator. Next, 10 μl of this mixture was loaded into the lanes of a precast 4-12% gradient polyacrylamide gel (4–12% Bolt Bis-Tris Plus Gels, Thermo Fisher Scientific). Molecular weights were determined using Precision Plus Protein Prestained Standards (Bio-Rad Laboratories). After electrophoresis, the proteins were transferred onto a 0.45-μm polyvinylidene difluoride membrane (Millipore). Next, after blocking for 30 min at room temperature and washing, the membrane was incubated with the primary antibody in Tris-buffered saline (20 mM Tris and 150 mM NaCl, pH = 7.4) supplemented with 0.1% Tween20 (TBS-T) containing 5% nonfat milk or Blocking One (Nacalai Tesque) overnight at 4°C. After washing with TBS-T, the membranes were incubated with horseradish peroxidase (HRP)-conjugated secondary antibody in TBS-T containing 5% nonfat milk or Blocking One solution at room temperature for 1 h. After washing with TBS-T, the membranes were treated with either the ECL start reagent (GE Healthcare) or the ECL select reagent (GE Healthcare) according to the manufacturer’s guidelines. The chemiluminescent images of the membranes were acquired using FUSION-SOLO.7S (Vilber-Loumat) and analyzed using ImageJ.

### Dot blot analysis

Lysates of PC3 cells and PC3 cell-derived sEVs were prepared as described above. The phospholipid content of these lysates was quantified using a LabAssay Phospholipid (Fujifilm). Subsequently, 100 μl of these lysates (phospholipid concentration = 6.4 μM) was adsorbed to a nitrocellulose membrane using a Bio-Dot (Bio-Rad). After blocking with TBS-T containing 3% BSA for 30 min at room temperature, followed by washing, the membrane was incubated with biotin-conjugated CTXB for the detection of GM1 (1:500, list labs, Cat# 112) or with the primary IgM antibody against GM3 clone GMR6 (1:1000, Tokyo Chemical Industry, Cat# A2582) in TBS-T containing 3% BSA overnight at 4°C. After washing with TBS-T, the membranes were incubated with the HRP-conjugated primary antibody against biotin (1:10000, Cell Signaling, Cat# 5571) or the HRP-conjugated secondary antibody against IgM (1:10000, Invitrogen, Cat# 31440) in TBS-T containing 3% BSA solution at room temperature for 1 h. After washing with TBS-T, the membranes were incubated with the ECL start reagent (GE Healthcare). Chemiluminescence images of the membranes were acquired using FUSION-SOLO.7S (Vilber-Loumat) and analyzed using ImageJ.

### Preparation of ganglioside-containing liposomes

The gangliosides GM1 (bovine brain), GD1a (bovine brain) and GD2 (human brain) were purchased from Avanti Polar Lipids, AdipoGen Life Sciences and Santa Cruz, respectively. GM2, GM3, and GD3 were prepared by total chemical synthesis (Koikeda et al., 2019; Takahashi et al., 2020). In an eggplant-shaped flask, 50 μl of 1 mM ganglioside (GM1, GM2, GM3, GD1a, GD2, or GD3) in methanol, 500 μl of 10 mM DMPC in chloroform, and 0.5 μl of 1 mM Bodipy-SM (Invitrogen) in methanol were mixed in a solution of 99:1:0.01 mol% ratio DMPC:ganglioside:Bodipy-SM. Since the molar ratio of GM1 to lipids in PC3 cells is known to be 0.0158 mol% (Llorente et al., 2013) and since the relative density of GM1 in the sEVs derived from PC3 was 29.7 ± 8.1 (mean ± S.E., n=3) times greater than that in PC3 cells (Fig. S4B). Since gangliosides are distributed in both the outer and inner leaflets of liposomal membranes, the density of GM1 in sEVs is similar to that in the outer leaflet of liposomes containing 0.94 mol% GM1. Therefore, we prepared liposomes containing 1 mol% ganglioside. The solvent was first dried with nitrogen gas and further dried with a CVE-3000 (EYELA) evaporator with VT-2000 (EYELA). Then, 5 ml of PBS was added to the flask, which was incubated at room temperature for 1 h. The solution was completely frozen in liquid nitrogen and then thawed in hot water (45 ℃). The freeze‒thaw process was repeated 5 times. Using an extruder (LiposoFAST, AVESTIN), 500 μl of the vesicle suspension was passed through 100-nm pore-sized polycarbonate membrane filters (10 times). The amount of sialic acid in the liposomes was determined using a Sialic Acid Assay Kit (MAK314-1KT, Sigma Aldrich), and the concentration of liposomes containing a ganglioside was adjusted to the same level for use in microscopic observation.

### Immunofluorescence imaging of the extracellular matrix

MRC-5 and HS-5 cells were cultured on glass-bottom dishes (IWAKI) for two days. The cells were then incubated with 10 μg/ml primary antibodies against fibronectin (Sigma‒Aldrich, F3648), laminin (Sigma‒Aldrich, L9393), collagen type I (Abcam, ab34710), laminin α1 (Santa-cruz biotech, sc-74418) or laminin α5 (Abcam, ab17107) in MEM at 37°C for 30 min. Next, the cells were washed twice with HBSS and incubated with 12 μg/ml rhodamine-conjugated anti-rabbit IgG antibody (Cappel, 55666) or 10 μg/ml FITC-conjugated anti-mouse IgG antibody (Cappell, 55514) in MEM at 37°C for 30 min. After washing twice with HBSS, fresh HBSS was added to the glass-base dishes. Subsequently, the cells were observed by epi-fluorescence microscopy (ECLIPSE Ts2-FL, Nikon, 60x 1.40 NA oil objective), which was performed with a CMOS camera (DS-Qi2, Nikon). A 560 nm LED lamp (Nikon) was used to excite rhodamine. The fluorescence intensities of laminin in the regions of cells (cell contours were determined using bright-field images) were quantified using ImageJ. The actual fluorescence intensities of laminin in the region of cells were measured by subtracting the average fluorescence intensity without the primary antibody from the fluorescence intensity of laminin stained with both the primary and secondary antibodies.

### Determination of the concentrations of sEVs containing fluorescently labeled marker proteins by total internal reflection fluorescence microscopy (TIRFM)

The concentrations of protein and lipids in the sEV suspension (on the order of nM) were too low to determine very quantitatively via conventional spectrophotometry. Therefore, we employed single-fluorescent particle tracking to directly quantify the number of sEVs. In brief, the glass windows of single-well or triple-well glass-base dishes (IWAKI) were coated with 100 or 50 μl of 10 μg/ml anti-CD63 IgG antibody (8A12, Cosmo Bio), anti-CD81 IgG antibody (JS-81, BD Biosciences), or anti-CD9 antibody (M-L13, BD Biosciences) in HBSS and then incubated for 2 h at 37°C. Subsequently, the antibody solution was removed, and the glass window was coated with either 100 or 50 μl of 50 μg/ml casein (Sigma‒Aldrich) in HBSS for 1 h at 37°C. This coating process was essential for mitigating the nonspecific binding of sEVs to the glass surface during subsequent experiments.

Either 100 or 50 μl of three distinct concentrations of sEVs containing CD63-Halo7, CD81-Halo7, and CD9-Halo7, all conjugated with SF650T in HBSS, was applied to glass windows pre-coated with antibodies against CD63, CD81, and CD9, respectively. The samples were then incubated at 37°C for 1 h. After removing the sEVs and washing twice with HBSS, the individual fluorescent particles of the sEVs were observed at 37°C with single-molecule detection sensitivity by total internal reflection fluorescence microscopy (TIRFM) using an Olympus IX-83 microscope (60x 1.49 NA oil objective) equipped with a high-speed gated image intensifier (C9016-02MLG; Hamamatsu Photonics) coupled to an sCMOS camera (ORCA-Flash4.0 V2; Hamamatsu Photonics), as previously described (Komura et al., 2016; Kinoshita et al., 2017; Morise et al., 2019). SF650T was excited using a 647-nm laser (LuxXPlus647-140, 140 mW, Omicron Laserrange) at an intensity of 0.3 μW/μm^2^.

All the acquired sEV movies were analyzed using ImageJ (Fiji). First, a noise reduction process was carried out, wherein 10 frames of sEV images observed at 30 frames/sec were averaged. Subsequently, each pixel was replaced by the average of the 3×3 neighborhood pixels using a smooth filter. Single particles of sEVs whose fluorescence intensity exceeded that of free SF650T were identified, and the numbers of sEVs were counted. By performing these observations at three distinct sEV concentrations, we obtained a calibration curve. Thereafter, sEVs derived from both intact cells and integrin subunit KO cells were prepared at identical concentrations (0.8-4×10^10^ particles/ml).

### Determination of the numbers of sEVs bound to ECM components on glass

To prevent nonspecific binding of sEVs to the glass surface, 50 or 100 μl of 50 μg/ml casein solution in HBSS was added to glass windows in single-well or triple-well glass-base dishes (IWAKI) and incubated for 1 h at 37°C. Then, the casein solution was replaced with 50 or 100 μl of 100 μg/ml fibronectin (Roche), 50 μg/ml laminin (Sigma‒Aldrich), 20 μg/ml laminin-511 (BioLamina), 100 μg/ml collagen type I (Sigma‒Aldrich), or 50 μg/ml casein (Sigma‒Aldrich) for 2 h at 37°C. After removal of the ECM or casein solution, 50 or 100 μl of sEVs containing CD63/CD81/CD9-Halo7 labeled with SF650T at the same concentration was applied to the glass and incubated for 1 h at 37°C. To assess the relative contributions of integrin and GM1 to sEV binding to laminin, we performed a competitive inhibition assay using the glycan moiety of GM1, synthesized as previously reported (Yanaka et al., 2019). sEVs were applied to laminin-coated glass in the presence of a high concentration (0.5 mM final) of the GM1 glycan moiety. After removal of the sEV suspension and two washes with HBSS, individual fluorescent particles of the sEVs in HBSS were visualized at 37°C with single-molecule detection sensitivity by TIRFM. The number of sEVs bound to the ECM or casein on glass was determined using ImageJ (Fiji) as described above. The number of sEVs bound to ECM-coated glass was estimated by subtracting the number of sEVs bound to casein-coated glass in a 1024 pixel × 1024 pixel (81.9 μm × 81.9 μm) image.

### Determination of the numbers of sEVs bound to cell membranes

sEVs containing CD63-Halo7 labeled with SF650T (sEV-CD63Halo7-SF650T) were incubated with MRC-5 or HS-5 cells expressing mCherry or GFP in the cytosol at 37°C. sEV-CD63Halo7-SF650T was visualized with single-molecule detection sensitivity by TIRFM, and mCherry or GFP was observed by oblique illumination. The number of sEVs on the cell membrane (the cell contour was determined using binarized mCherry or GFP images) was quantified every 10 min by ImageJ (Fiji) as described above.

sEVs bound to cell membranes were visualized using confocal fluorescence microscopy. sEVs-CD63Halo7-TMR were incubated with MRC-5 or HS-5 cells expressing GFP at 37℃ for 1 hour. After incubation, the cells were fixed with 4 % paraformaldehyde (PFA) at room temperature for 30 min. After removing the PFA solution, the cells were treated with 0.1 M glycine for 5 min to quench residual fixation. The samples were then washed with HBSS, and an aqueous mounting medium (Thermo Fisher Scientific) was applied. Fluorescent images of sEVs-CD63Halo7-TMR and GFP within the cells were acquired using an FV-1000 confocal microscope (Olympus) with a prolonged detector exposure period. The number of sEVs associated with cell membranes was quantified as described above.

### Observation of sEVs stained with antibodies targeting activated integrin β1

sEVs containing CD63-Halo7 labeled with TMR (sEV-CD63Halo7-TMR) were incubated on either uncoated glass or laminin-511-coated glass for 1 hour. After incubation, the samples were washed with HBSS, and 10 μg/ml of general anti-integrin β1 antibody (clone P5D2 from Abcam), or antibodies specific for activated integrin β1 (clone HUTS-4 from Millipore and clone HUTS-21 from BD Pharmingen) (Luque et al., 1996), was added and incubated at 37℃ for 30 min. After two washes with HBSS, 1 μg/ml AlexaFlour488-conjugated anti-mouse IgG antibody was applied and incubated at 37℃ for 30 min. The glass was then washed five times with HBSS. Single fluorescent particles of sEV-CD63Halo7-TMR and AlexaFlour488-conjugated antibody on uncoated glass or laminin 511-coated glass were simultaneously observed by TIRFM. Colocalization between sEV-CD63Halo7-TMR and anti-integrin β1 antibody was analyzed using ImageJ (Fiji). The brightest point of each sEV-CD63Halo7-TMR signal was identified as the center of the sEV. The fluorescence intensity of AlexaFlour488 within a circular region with a radius of 3 pixels, centered on the detected coordinates of the sEV, was quantified. Instances where the AlexaFlour488 intensity exceeded the signal of a single AlexaFlour488 molecule, were recorded as colocalization events. Finally, the percentage of colocalization events relative to the total number of sEVs was calculated, representing the ratio of sEVs stained with anti-integrin β1 antibody.

### Simultaneous dual-color observation of dSTORM movie of ECM components and single fluorescent particles of sEVs on living cell plasma membranes

For immunostaining of the ECM components for dSTORM movie observation, 0.67 μl of 5.6 μg/ml SaraFluor650B (SF650B, Goryo Chemical) NHS ester was incubated with 100 μl of 1 mg/ml anti-rabbit IgG antibody (0212-0081, Cappel) in 0.1 M NaHCO_3_ for 60 min at room temperature. SF650B-conjugated anti-rabbit IgG was then isolated using a Sephadex G-25 column (GE Healthcare). Since SF650B spontaneously blinks, it enables the observation of dSTORM movie observations in living cell PMs. MRC-5 cells cultured on glass-based dishes for 2 days were subsequently incubated with 10 μg/ml anti-fibronectin IgG (F3648; Sigma‒Aldrich), anti-laminin IgG (L9393; Sigma‒Aldrich) or anti-collagen type I IgG (ab34710; Abcam) for 30 min at 37°C. After washing twice with HBSS, the cells were incubated with 10 μg/ml SF650B-conjugated anti-rabbit IgG antibody for 30 min at 37°C. After washing with HBSS 3 times, sEVs containing CD63-Halo7-TMR were incubated with the cells for another 30 min at 37°C. Individual fluorescent particles of the sEVs and single molecules of SF650B on the apical surface of the cells at 512×512×pixel (25.6 μ×25.6 μm) were observed at 5-ms resolution (200 frames/second) for 3504 frames by oblique-angle illumination using an Olympus IX-83 inverted microscope (100×1.5 NA oil objective) equipped with two high-speed gated image intensifiers (C9016-02MLG; Hamamatsu Photonics) coupled to two sCMOS cameras (ORCA-Flash4.0 V2; Hamamatsu Photonics), as described previously (Komura et al., 2016; Kinoshita et al., 2017; Morise et al., 2019; Kemmoku et al., 2024). TMR and SF650B were excited using a 561-nm laser (Excelsior-561-100-CDRW, 100 mW, Spectra-Physics) and a 647-nm laser (140 mW, Omicron Laserrange) at 2.8 and 16 μW/μm^2^, respectively. The final magnification was 133×, yielding pixel sizes of 47.1 nm (square pixels). To obtain a single dSTORM image, data acquisition was performed for 1002 frames and then repeated by shifting the initial frames backward by 6 frames, resulting in a total of 417 dSTORM images (Fig. 6A). By connecting these dSTORM image sequences, we generated a pseudo-real-time “dSTORM movie”. Images of individual sEV particles were concurrently recorded at 200 frames/second, and images averaged over 6 frames were subsequently combined to produce the movie. The pseudo-real-time movies of ECM components and sEV particles (33 frames/second) were superimposed (Fig. 6A).

More details regarding the data acquisition process for dSTORM imaging and the generation of the dSTORM movie are provided below. The dSTORM super-resolution video data were generated using frame information and the x and y coordinates of all the spots, which were incorporated into the CSV data output by the ThunderSTORM plugin for ImageJ (Ovesný et al., 2014) installed in the Fiji package (Schindelin et al., 2012). dSTORM image reconstructions were conducted with a pixel size of 10 nm. Gaussian rendering with a localization precision of 24 nm was utilized in this process. Furthermore, we used the uncertainty that represents the spatial precision of the spot. A Gaussian distribution was generated for each spot, with the x and y coordinates serving as the center and the standard deviation (S.D.) being equivalent to 6 times the uncertainty of the spot. This distribution indicates the existence probability of the spot. The existence probability distribution for all structures at time t was obtained as a summation of all the distributions for spots that appeared in [6t+1 6t+1002] frames. The dSTORM super-resolution video data were generated by repeating this process for time t=0,1,2,… If the value of each time and coordinate in the dSTORM video data exceeded a threshold value θ, we considered a structure to exist at the time and coordinate. The optimal threshold value θ was determined for each dSTORM video data point to minimize the average in-class variance for the two classes when every value in the dSTORM video data was classified into two classes using the threshold value θ (Chen et al., 2014; Kemmoku et al., 2024). To synchronize the dSTORM movie of ECM structures with the single-particle movie of sEVs, the temporal resolution of the single-particle movie was converted to 33.3 frames/s by averaging 6 frames, and the averaged single-particle image was merged with the dSTORM image created from 1002 frames, of which the middle frame was synchronized with the averaged single-particle image (Fig. 6A). Single-particle tracking in the movie was performed by in-house computer software based on previously reported methods (Gelles et al., 1988; Kusumi et al., 1993). More details have been described previously (Suzuki et al., 2007a, 2007b; Kasai et al., 2011; Suzuki et al., 2012; Komura et al., 2016; Kinoshita et al., 2017; Morise et al., 2019; Fujiwara et al., 2023a, 2023b).

### Colocalization analysis between ECM structures in dSTORM movies and single sEV particles

The distances between the centroids of the sEVs and the boundaries of the ECM structures were quantified using single-particle tracking data and binarized dSTORM movies. To generate binarized images of the ECM structures in the dSTORM movie, we employed the kernel density estimation (KDE) method (Kemmoku et al., 2024). KDE is a traditional image segmentation method that can rapidly determine the criterion for segmenting PALM and dSTORM images, as reported previously (Chen et al., 2014; Kemmoku et al., 2024). Furthermore, a recent study demonstrated that KDE is one of the most appropriate methods for segmenting images of clearly outlined structures, such as protein clusters (Nieves et al., 2023). KDE provides a way to interpolate object boundaries without bias by using random fluctuations of activated boundary fluorophores.

The ThunderSTORM datasets were imported into in-house KDE software (MATLAB) for image segmentation and colocalization analysis. We used not only the localization coordinates but also the uncertainty, which reflects the spatial detection precision of the spots. Let *x*_*i*_, *y*_*i*_ and *u*_*i*_, *i* = 1,2, ⋯, *N* be the horizontal and vertical localization coordinates and the uncertainty of spot *i*, respectively. *N* represents the total number of spots. Considering the uncertainty of the location measurement of spot *i*, the existence probability *p*_*i*_(*x*, *y*) of spot *i* is considered to spread to the Gaussian distribution with the center coordinate *x*_*i*_, *y*_*i*_ and the standard deviation (S.D.) that is proportional to the uncertainty *u*_*i*_ as follows:

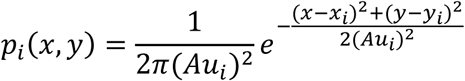

in which *A* is the proportional coefficient of S.D. to the uncertainty *u*_*i*_. This coefficient was estimated as *A* ≈ 6 by some preliminary experiments. The Gaussian distribution for constructing each existence probability *p*_*i*_(*x*, *y*) is called the Gaussian kernel. The existence probability distribution for all spots, i.e., the dSTORM image, is obtained by averaging the existence probability distribution *p*_*i*_(*x*, *y*) as follows:

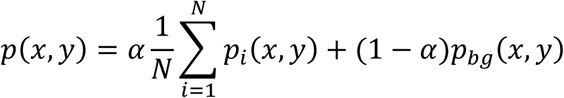

excluding spots having extremely small or large uncertainties *u*_*i*_ because such spots are likely to contain noise (e.g. electrical shot noise from camera). *p_*b*g_*(*x*, *y*) represents the background noise generated by the probability density distribution 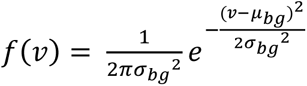, independent of *x*, *y*. *⍺* is a weight determined by the respective occurrence probabilities of the spots and background noise.

The structures can be considered to exist where the dSTORM image, which represents the probability of structure existence, is above a certain threshold value *θ* because the spots appear uniformly inside the structures with a certain probability. It is necessary to determine the optimal threshold value *θ* for each obtained dSTORM image because the optimal value differs depending on the target molecule and the experimental environment. In general, the histogram of *p*(*x*, *y*) values for all *x*, *y* forms a bimodal shape consisting of both larger values created by dense spots appearing in structures and smaller values created by sporadic noise. Therefore, the threshold value *θ* should be determined to separate these two structures. Otsu’s method is a well-known approach for determining such a threshold value (Otsu, 1979). Let *S*_*in*_ and *S*_*out*_ be sets of coordinates (*x*, *y*) where *p*(*x*, *y*) ≥ *θ* and *p*(*x*, *y*) < *θ*, respectively. The sets *S*_*in*_ and *S*_*out*_ indicate the inside and outside of the clusters, respectively. The numbers of elements in sets *S*_*in*_ and *S*_*out*_ are represented by *N*_*in*_ and *N*_*out*_, respectively. The intraclass variances of the sets *S*_*in*_ and *S*_*out*_ are calculated by

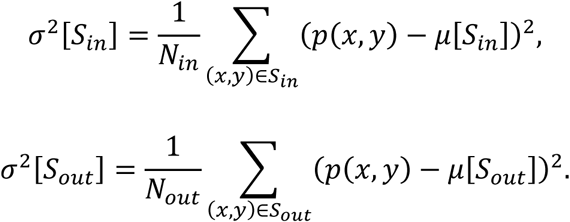

In the above equations, *μ*[*S*_*in*_] and *μ*[*S*_*out*_] are the respective averages of the set *S*_*in*_ and *S*_*out*_ obtained by

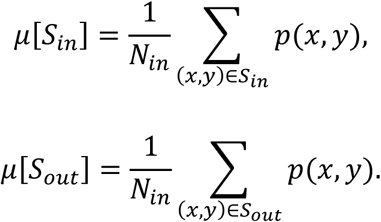

The average of these intraclass variances is

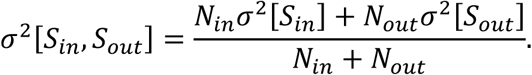

In Otsu’s method, the optimal threshold value *θ̂* to minimize the average of the intraclass variances *σ*^2^[*S*_*in*_, *S*_*out*_] is determined as follows:

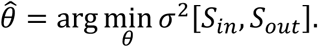

In this study, the structures were determined using Otsu’s method.

Thereafter, the distance from the contour of the determined structures to the center of the sEVs on the PM was measured in all the frames of the superimposed movies of the ECM components and sEVs, and the number densities of pairwise distances at a given distance were plotted. If an sEV particle was within an ECM structure, the measured distance was expressed as a negative value. Furthermore, we simulated the distances between random coordinates and the boundaries of ECM structures in silico and calculated their corresponding relative frequency distributions using a bin width of 50 nm. To normalize the relative frequency of sEVs within each bin, we divided the data by the corresponding relative frequency of random spots. A ratio exceeding 1 indicated a higher incidence of sEVs at the measured distance.

### dSTORM imaging of ECM-coated glass

The ECM molecules were coated on triple-well glass-bottom dishes and incubated with 5 μg/ml anti-fibronectin IgG (F3648; Sigma‒Aldrich), anti-laminin IgG (L9393; Sigma‒Aldrich), or anti-collagen type I IgG (ab34710; Abcam) for 2 h at room temperature. After washing twice with HBSS, the ECM molecules were incubated with 2.5 μg/ml SF650B-conjugated anti-rabbit IgG antibodies for 1 h at room temperature. After washing with HBSS 5 times, single molecules of SF650B at 512 pixels × 512 pixels (25.6 μm × 25.6 μm) were observed via TIRFM at 5-ms resolution (200 frames/s) for 1008 frames. The dSTORM images were reconstructed from the movie using the ThunderSTORM plugin for ImageJ.

### Cell spreading assay

The cells that secreted sEVs were cultured for 3-5 days and subsequently collected using a solution of 0.5% BSA and 5 mM EDTA in PBS. A total of 4×10^5^ cells/ml were then evenly dispersed onto an ECM-coated single-well glass-bottom dish. The cells were observed after incubation for 0, 30, 60, or 120 min with an epi-illumination microscope (CKX53, Olympus, 20x 0.40 NA objective) equipped with a camera (WAT-01U2, WATEC). The cell area was quantified using ImageJ software.

### Observation of branched protrusions in HUVEC after sEV treatment

HUVECs were transfected with a plasmid encoding GFP and siRNA targeting laminin γ1 (SI02757475, Qiagen). After three days of culture, the transfected HUVEC were seeded onto fibronectin-coated glass-bottom dishes and cultured for an additional day. Subsequently, the cells were incubated with PC3-derived sEVs or 0.26 nM (10 ng/ml) VEGF (Peprotech) for 12 hours. The HUVEC-GFP and HUVEC-GFP-LNγ1KD were then visualized using TIRFM. The position of the greater curvature on the HUVEC membrane was designated as the origin of the branched protrusion. The length of the protrusion, defined as the distance between the branched protrusion origin and the tip of the protrusion (Fig. 9F), was measured following the previously reported protocol (Myers et al., 2011; Braun et al., 2014).

## Supplemental material

Fig. S1 shows the preparation of sEVs derived from PC3 cells and the determination of their size and concentration. Fig. S2 shows the number of sEV particles attached to glass coated with ECM components. sEVs containing CD81 or CD9-Halo7 were derived from wild-type, integrin β1, integrin α2, integrin α6, or integrin β4 KO PC3 cells and labeled with SF650T. Fig. S3 shows the results of western blot analysis to determine whether sEVs derived from PC3 cells were covered by fibronectin and/or laminin. Fig. S4 shows the number of sEVs derived from B78 cells expressing several glycosyltransferases and the number of liposomes containing gangliosides bound to laminin. Fig. S5 shows the flow cytometry analysis of B78 cells expressing several glycosyltransferases to examine the expression levels of gangliosides. Fig. S6 shows the number of PC3-derived sEVs bound to laminin on glass and MRC-5 PMs before and after cholesterol depletion or addition. Table S1 presents the numbers of sEVs labeled with tetraspanin-Halo7-TMR attached to glass coated with ECM components. Table S1 is related to Figs. 3 and S2. Videos 1-3 (Video 1: fibronectin, Video 2: collagen type I, Video 3: laminin) show the simultaneous observation of single sEV-CD63Halo7-TMR particles (green) and dSTORM ECM movies (magenta) on an MRC cell. Video 4 shows an enlarged movie of the colocalization of an sEV-CD63Halo7-TMR particle (green) and laminin (dSTORM movie, magenta) on an MRC cell.

## Data availability

The data supporting the findings of this study are available from the corresponding author upon reasonable request.

## Acknowledgments

We thank Joan Massagué (Memorial Sloan Kettering Cancer Center) and Ayuko Hoshino (University of Tokyo) for kindly providing the human 4175-LuT cells. We also thank Yumi Matsuno for supporting the preparation of extracellular vesicles from tumor cells, Shinobu Kawaguchi for constructing various cDNAs, and Takahiro Fujiwara and Akihiro Kusumi for developing the analysis software for single-molecule imaging. This work was supported in part by Japan Science and Technology Agency (JST) grants from the Core Research for Evolutional Science and Technology (CREST) program in the field of “Extracellular Fine Particles” (JPMJCR18H2) (K.G.N.S., K.M.H., and H.A.) and “Cell Control” (JPMJCR24B3) (K.G.N.S., K.M.H.), Grants-in-Aid for Scientific Research from the Japan Society for the Promotion of Science (JSPS) (21H02424, 18H02401) (K.G.N.S.), a Grant-in-Aid for Challenging Research (Exploratory) from JSPS (20K21387) (K.G.N.S.), a Grant-in-Aid for Core-to-Core Program from JSPS (JSCCA202000007) (K.G.N.S., and H.A.), a Grant-in-Aid for Innovative Areas from the Ministry of Education, Culture, Sports, Science and Technology of Japan (MEXT) (18H04671) (K.G.N.S.), the National Cancer Center Research and Development Fund (2023-A-03) (K.G.N.S.), the Takeda Science Foundation (K.G.N.S.), Uehara Memorial Foundation (K.G.N.S.), the Mizutani Foundation for Glycoscience (K.G.N.S.), the Daiichi Sankyo Foundation of Life Science (K.G.N.S.), the Research Foundation for Opto-Science and Technology (K.G.N.S.), and the Naito Foundation (K.G.N.S.), as well as a Grant-in-Aid from Support for Pioneering Research Initiated by the Next Generation in JST (JST SPRING) (JPMJSP2125) (T. I.) and a Grant-in-Aid for JSPS Fellows (23KJ1045) (T. I.).

## Author contributions

T.I., K.M.H., and K.G.N.S. conceived and formulated the project. T.I., K.M.H., R.S.K. and K.G.N.S. designed the biological experiments and participated in the discussions. T.I., M.K., and A.S. performed the experiments on the binding of EVs to ECM components and the data analysis. Y.M. helped to isolate sEVs from cell culture supernatants. S.K. generated the cDNA plasmids. Koichi F, Y.O., and Keiko F. generated B78 cells expressing only one type of ganglioside. H.A. and N.K. synthesized the gangliosides. Y.Y., T.I., K.M.H., and K.G.N.S. developed the analysis software for super-resolution movies and single-particle tracking. T.I., Y.Y. and K.G.N.S. wrote the manuscript, and all the authors discussed the results and participated in revising the manuscript.

## Author notes

Disclosures: The authors declare no competing interests exist.

