## Supplementary Material for "Extracellular vesicles adhere to cells primarily by interactions of integrins and GM1 with laminin"

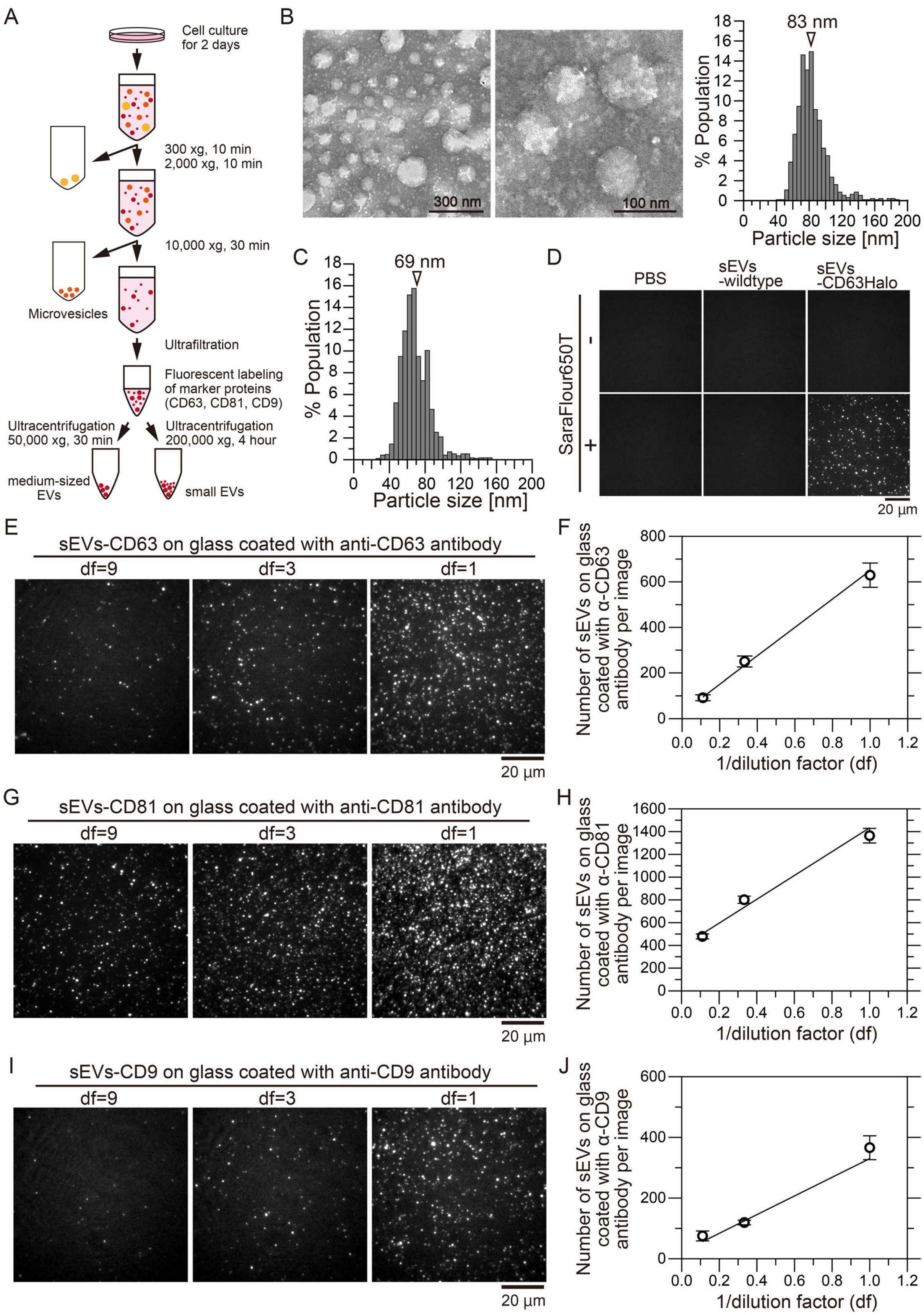
A
Cell culture for 2 days
300 xg, 10 min
2,000 xg, 10 min
10,000 xg, 30 min
Microvesicles
Ultrafiltration
Fluorescent labeling of marker proteins (CD63, CD81, CD9)
Ultracentrifugation 50,000 xg, 30 min
Ultracentrifugation 200,000 xg, 4 hour
medium-sized EVs
small EVs
B
300 nm
100 nm
83 nm
% Population
Particle size [nm]
C
69 nm
% Population
Particle size [nm]
D
PBS
sEVs -wildtype
sEVs -CD63Halo
SaraFlour650T
-
+
20 µm
E
sEVs-CD63 on glass coated with anti-CD63 antibody
df=9
df=3
df=1
20 µm
F
Number of sEVs on glass coated with α-CD63 antibody per image
1/dilution factor (df)
G
sEVs-CD81 on glass coated with anti-CD81 antibody
df=9
df=3
df=1
20 µm
H
Number of sEVs on glass coated with α-CD81 antibody per image
1/dilution factor (df)
I
sEVs-CD9 on glass coated with anti-CD9 antibody
df=9
df=3
df=1
20 µm
J
Number of sEVs on glass coated with α-CD9 antibody per image
1/dilution factor (df)

**Fig. S1.** Preparation of sEVs derived from PC3 cells and determination of their size and concentration. (A) sEVs from PC3 cells were isolated from the cell culture supernatant by ultrafiltration and ultracentrifugation. Tetraspanins tagged with Halo7 in sEVs were fluorescently labeled with SaraFluor650T (SF650T). (B) Negative-staining TEM images of sEVs revealed that the mean size of the sEVs was 83 ± 19 nm (mean ± SD). (C) The mean size of the sEVs determined by qNano was 69 ± 17 nm, as indicated by the arrowhead. (D) Single-particle fluorescence images of sEVs by total internal reflection fluorescence microscopy (TIRFM). Only when sEVs expressed CD63-Halo7, single particles labeled with SF650T (sEVs-CD63Halo7-SF650T) could be observed. (E, G, and I) Single-particle fluorescence images of three concentrations of sEV-CD63Halo7-SF650T particles (E), sEV-CD81Halo7-SF650T particles (G), and sEV-CD9Halo7-SF650T particles (I), which attached to glass coated with anti-CD63 antibody, anti-CD81 antibody, and anti-CD9 antibody, respectively. df indicates the dilution factor. (F, H, and J) The numbers of sEV-CD63Halo7-SF650T particles (F), sEV-CD81Halo7-SF650T particles (H), and sEV-CD9Halo7-SF650T particles (J) at three df values attached to the CD63 antibody, anti-CD81 antibody, and anti-CD9 antibody-coated glass, respectively ($n$ = 16 images). Data are presented as the mean ± SE. The sEV concentration was adjusted according to the calibration line.

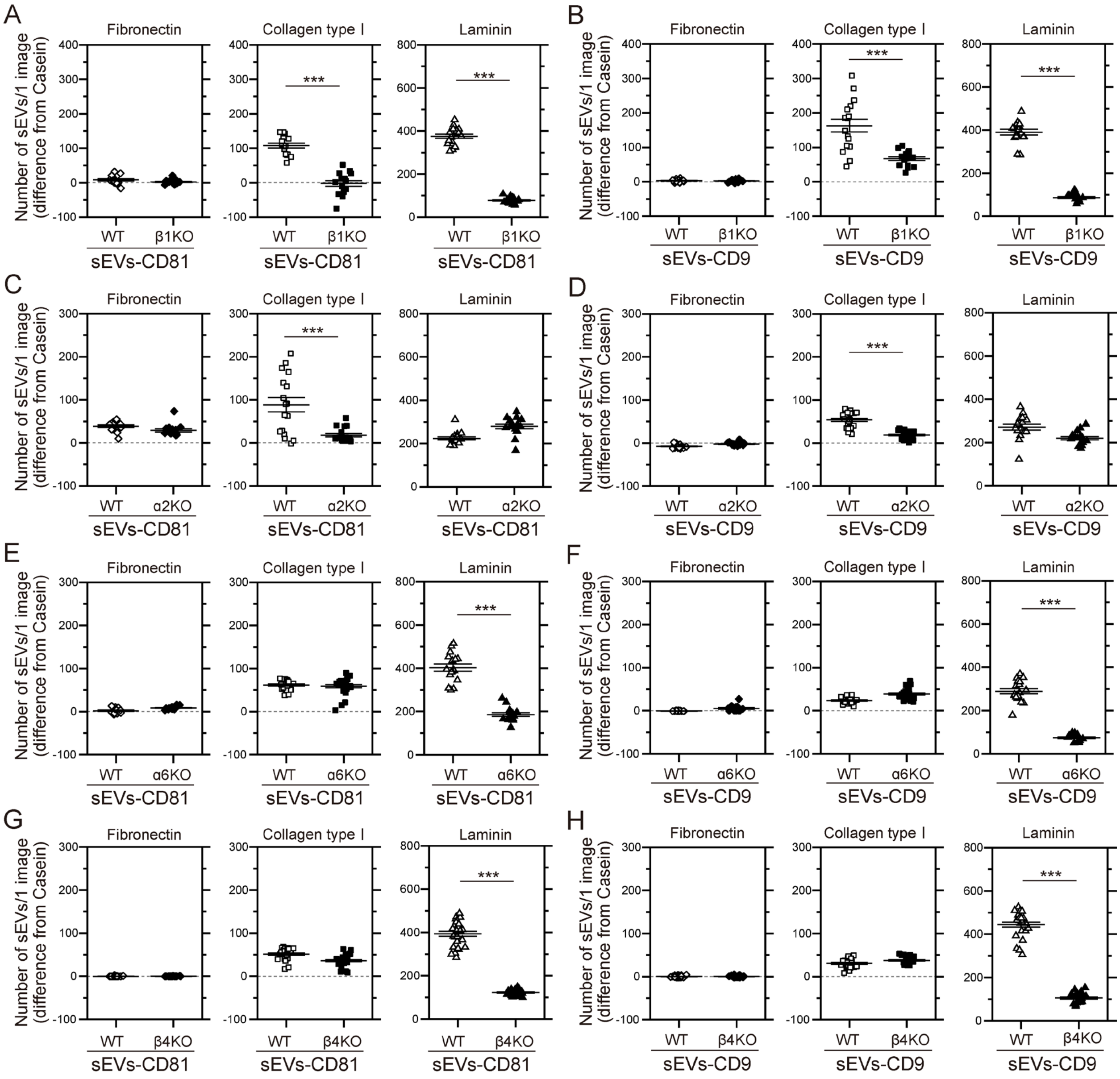


**Fig. S2.** The integrin β1, integrin α6 and integrin β4 subunits in sEVs containing CD81 or CD9 derived from PC3 cells are responsible for the binding of the sEVs to laminin, and the integrin α2 subunit is responsible for the binding of the sEVs to collagen type I. The numbers of intact PC3 cell-derived sEVs attached to glass coated with fibronectin, collagen I and laminin were compared with those of sEVs derived from integrin β1 (A and B), integrin α2 (C and D), integrin α6 (E and F) and integrin β4 (G and H) KO cells. CD81-Halo7 (A, C, E, and G) and CD9-Halo7 (B, D, F, and H) in sEVs were labeled with SF650T. Data are presented as the mean ± SE. n.s., non-significant difference; **P*<0.05; ***P*<0.01; ****P*<0.001 according to Welch's t-test (two-sided).

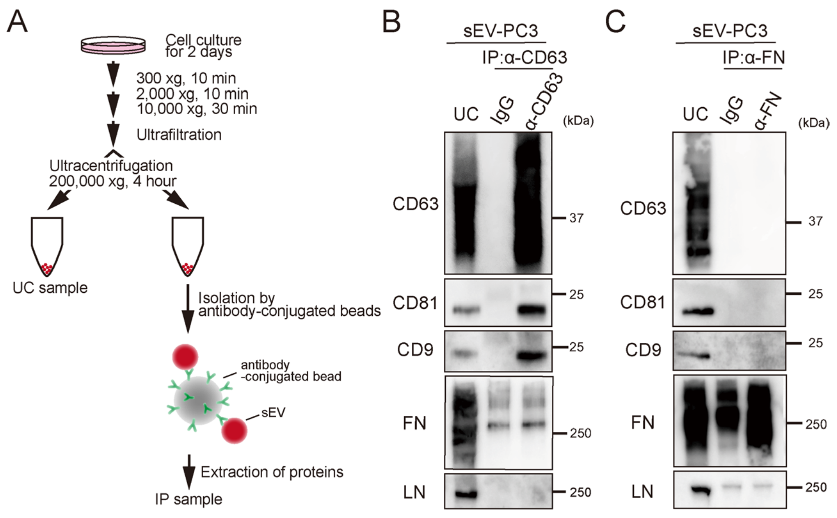


**Fig S3.** (A) Schematic diagram of the isolation of specific sEVs. sEVs were isolated by ultracentrifugation at 200,000 × g for 4 h (UC sample). Then, special sEVs were isolated from sEVs by a bead-conjugating antibody (IP: immunoprecipitation sample). (B) Western blot analysis of tetraspanin (CD63, CD81, and CD9), fibronectin (FN), and laminin (LN) in UC and IP samples. sEVs were isolated by immunoprecipitation using an anti-CD63 antibody. (C) Western blot analysis of sEVs isolated by immunoprecipitation using anti-fibronectin antibody.

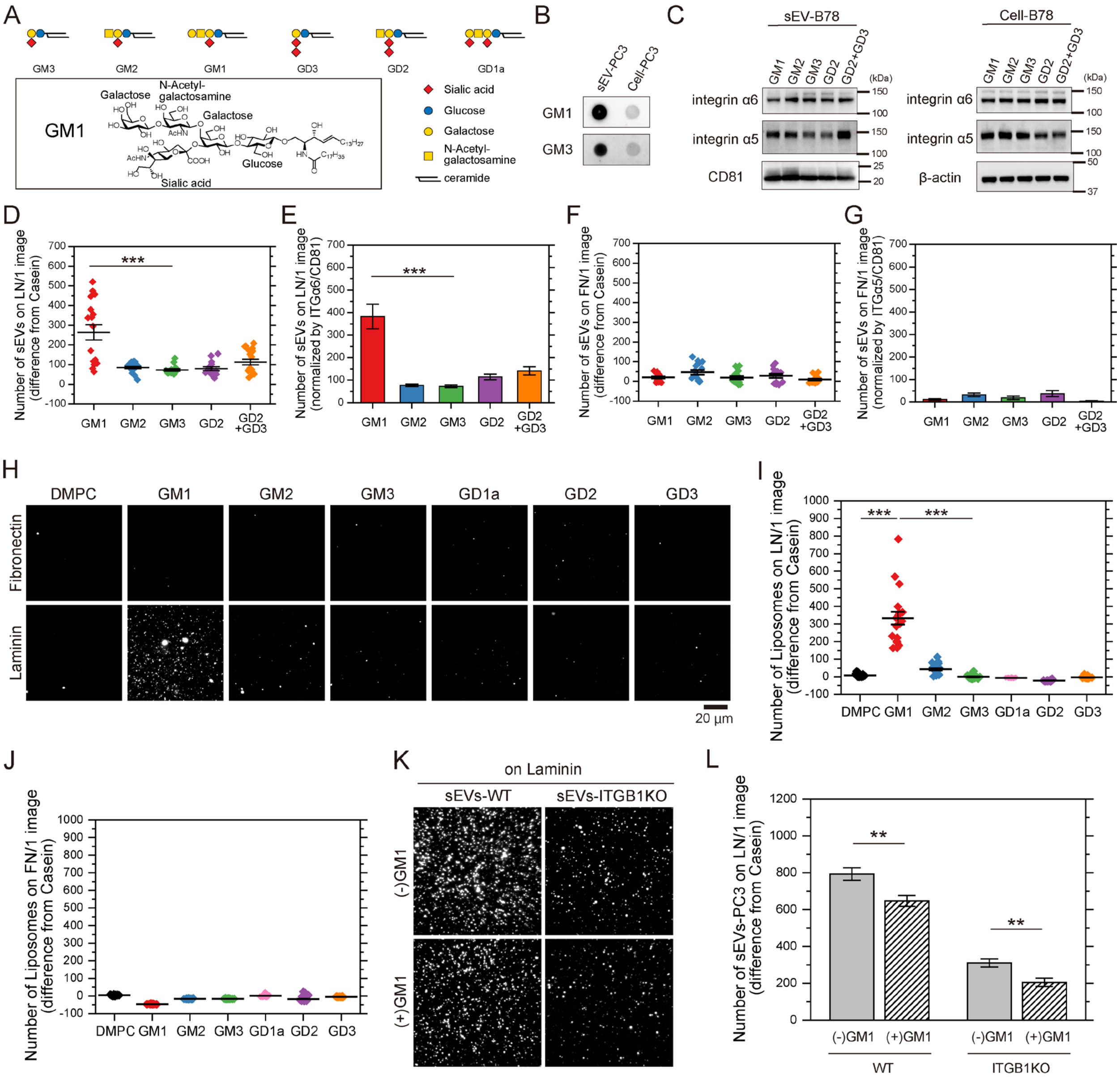


**Fig. S4.** GM1 is responsible for the binding of sEVs to laminin. (A) (top) Schematic diagram of gangliosides and (bottom) chemical structure of GM1. Monosaccharide symbols follow the Symbol Nomenclature for Glycans (SNFG). (B) Dot blotting of GM1 and GM3 in PC3 cells and PC3-derived sEVs. (C) Western blot analysis of integrin subunits in B78 cell lines with abundant expression of one type of ganglioside—GM1, GM2, GM3, GD2 or GD2/GD3—and sEVs derived from these cells. (D and E) The numbers of sEVs attached to glass coated with laminin (D) and the numbers normalized to the ratio of integrin α6/CD81 in the sEVs (E). (F and G) The numbers of sEVs attached to glass coated with fibronectin (F) and the numbers normalized to the ratio of integrin α5/CD81 in the sEVs (G). (H) Single-particle fluorescence images of DMPC-liposomes containing GM1, GM2, GM3, GD1a, GD2, or GD3 on glass coated with fibronectin or laminin. (I and J) The numbers of liposomes attached to glass coated with laminin (I) or fibronectin (J). (K) Single-particle fluorescence images of sEVs-PC3-CD63Halo7-TMR and sEVs-PC3-ITGB1KO-CD63Halo7-TMR on laminin before or after treatment of the GM1's glycan. (L) The numbers of sEVs bound to laminin before or after treatment with a high concentration of the GM1 glycan moiety (0.5 mM final). Data are presented as the mean ± SE. n.s., non-significant difference; **P*<0.05; ***P*<0.01; ****P*<0.001 according to Welch's t-test (two-sided).

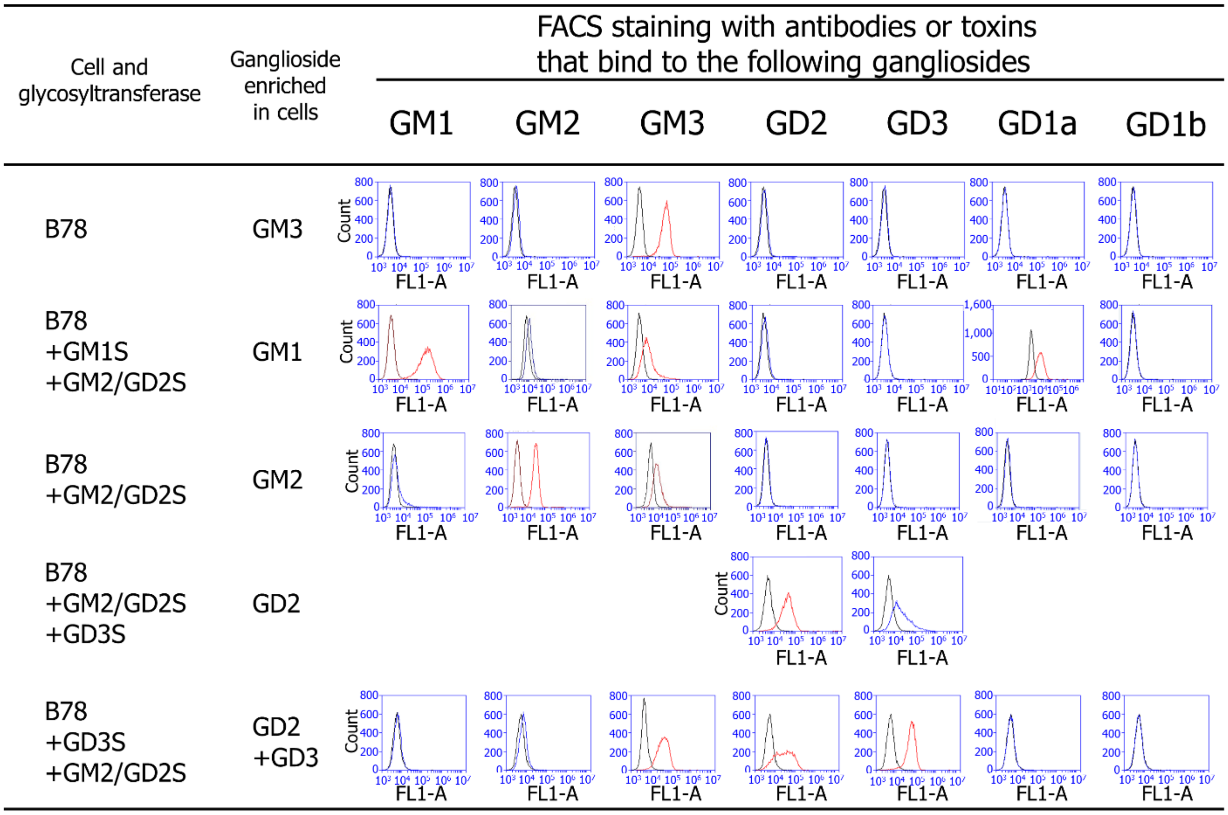


**Fig. S5.** Expression levels of gangliosides in B78 cell PMs analyzed by flow cytometry.

The cells were stained as described in the Materials and Methods. Control specimens were prepared without the primary antibodies or biotinylated cholera toxin B (black traces). The red traces indicate the molecules that were stained with antibodies (positive), and the blue traces indicate molecules that were not stained (negative) or were only slightly stained (weakly positive).

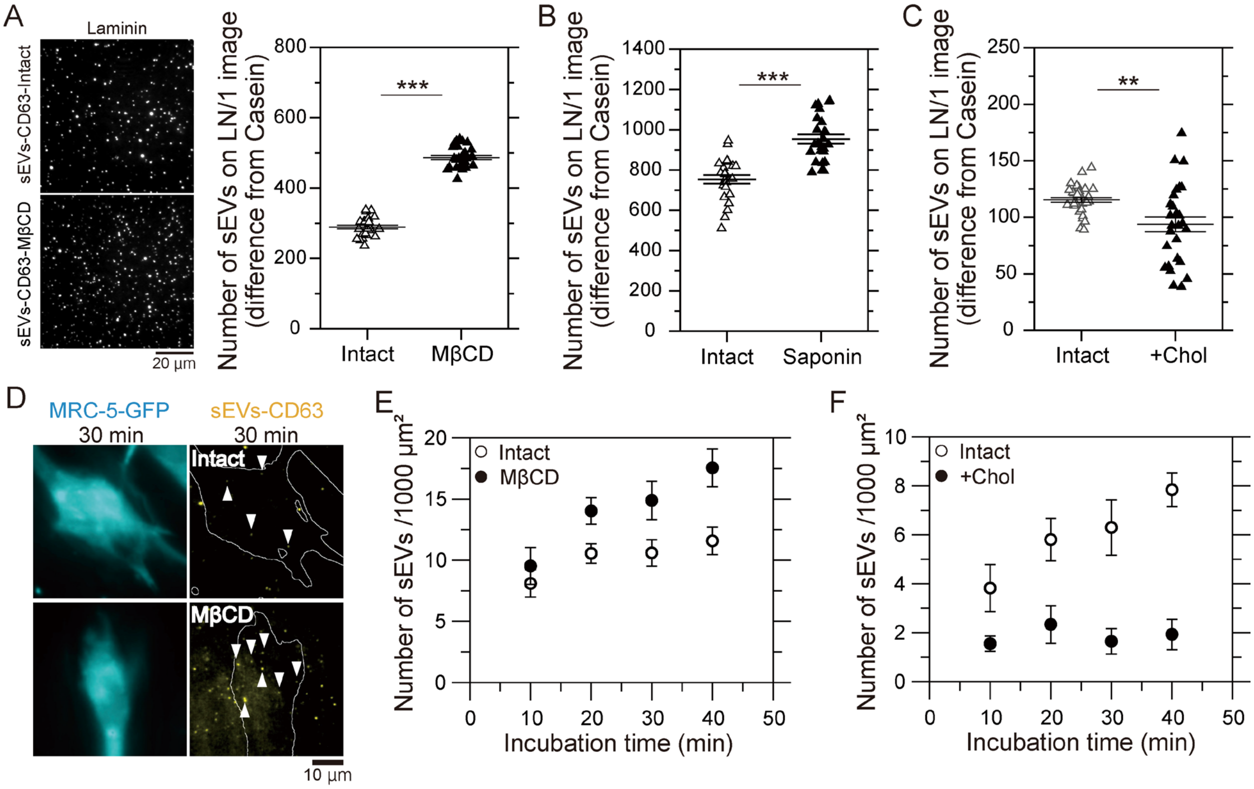


**Fig. S6.** (A) Fluorescence images of sEV-CD63Halo7-SF650T particles bound to laminin (LN) on glass before and after cholesterol depletion by MβCD and the numbers of attached sEVs per image (82 μm × 82 μm). The cholesterol content in PC3 cell-derived sEVs was reduced to 16% after treatment with MβCD. (B and C) The numbers of sEV-CD63-Halo7-SF650T particles attached to glass coated with laminin before and after treatment with saponin (B) and the addition of cholesterol by MβCD-cholesterol complex (C).The cholesterol content was increased to 186% after treatment with the MβCD-cholesterol complex. (D) Fluorescence images of an MRC-5-GFP cell and sEV-CD63Halo7-SF650T particles on the MRC-5 cell after 30 min of incubation. (E and F) Time course of the number of sEV-CD63-Halo7-SF650T particles per 1000

$\mu m^2$ attached to the MRC-5 cell membrane before and after treatment with MβCD ($n$ = 16 cells) (E) or the MβCD-cholesterol complex ($n$ = 8 cells) (F). Data are presented as the mean ± SE. n.s., non-significant difference; *$P$<0.05; **$P$<0.01; ***$P$<0.001 according to Welch's t-test (two-sided).

**Supplemental Table 1.** The numbers of sEVs labeled with tetraspanin-Halo7-TMR attached to glass coated with the ECM.

| sEV | Integrin | Fibronectin | | | Collagen type I | | | Laminin | | |
|---|---|---|---|---|---|---|---|---|---|---|
| | | Mean±SE | KO/WT* | p value† | Mean±SE | KO/WT | p value | Mean±SE | KO/WT | p value |
| CD63 | WT | 19±14.8 | | | 42±2.1 | 0.16±0.024 | $8.7\times10^{-16}$ | 424±10.5 | 0.22±0.008 | $1.2\times10^{-21}$ |
| | β1KO | 15±3.5 | | | 7±1.0 | | | 93±2.7 | | |
| | WT | 9±1.6 | | | 126±9.4 | 0.38±0.056 | $8.4\times10^{-7}$ | 437±21.1 | | |
| | α2KO | 20±4.7 | | | 48±6.0 | | | 380±8.6 | | |
| | WT | 10±1.0 | | | 66±1.3 | | | 220±6.0 | 0.24±0.011 | $9.9\times10^{-16}$ |
| | α6KO | 19±1.0 | | | 76±1.6 | | | 52±2.0 | | |
| | WT | 22±1.6 | | | 65±1.3 | | | 232±6.7 | 0.40±0.014 | $9.6\times10^{-16}$ |
| | β4KO | 9±0.8 | | | 73±1.5 | | | 93±1.8 | | |
| CD81 | WT | 8±2.9 | | | 108±7.0 | -0.02±0.076 | $1.8\times10^{-10}$ | 376±9.1 | 0.21±0.009 | $4.4\times10^{-19}$ |
| | β1KO | 2±1.6 | | | -2±8.2 | | | 77±2.8 | | |
| | WT | 39±2.7 | | | 88±16.2 | 0.20±0.057 | $6.9\times10^{-4}$ | 223±6.9 | | |
| | α2KO | 29±3.7 | | | 18±3.8 | | | 279±10.3 | | |
| | WT | 2±1.6 | | | 62±2.1 | | | 403±16.4 | 0.46±0.027 | $1.0\times10^{-10}$ |
| | α6KO | 9±0.8 | | | 59±3.7 | | | 186±7.9 | | |
| | WT | 0±0.1 | | | 51±2.5 | | | 393±11.4 | 0.31±0.011 | $3.2\times10^{-19}$ |
| | β4KO | 0±0.1 | | | 36±2.5 | | | 122±2.3 | | |
| CD9 | WT | 3±0.9 | | | 163±18.2 | 0.41±0.057 | $1.3\times10^{-4}$ | 390±12.4 | 0.22±0.014 | $4.5\times10^{-15}$ |
| | β1KO | 3±0.9 | | | 68±5.3 | | | 87±4.6 | | |
| | WT | -8±0.8 | | | 54±3.3 | 0.35±0.039 | $1.0\times10^{-10}$ | 271±13.5 | | |
| | α2KO | -2±0.7 | | | 19±1.7 | | | 220±7.3 | | |
| | WT | -1±0.2 | | | 23±1.2 | | | 289±11.2 | 0.26±0.017 | $4.3\times10^{-14}$ |
| | α6KO | 5±1.6 | | | 39±2.4 | | | 74±4.0 | | |
| | WT | 0±0.3 | | | 30±1.9 | | | 444±11.5 | 0.24±0.012 | $4.8\times10^{-24}$ |
| | β4KO | 0±0.4 | | | 37±2.0 | | | 106±4.4 | | |

*The ratio of the number of integrin KO PC3-derived sEVs bound to ECM components to the number of wild-type (WT) PC3-derived sEVs bound to ECM components is shown only when the ratio was less than 0.5 and the difference was greater than 20.

[†]p values of Welch's t-test (two-sided) are shown.

**Supplemental Videos 1-3.** Movies showing the simultaneous observation of sEV-CD63Halo7-TMR particles (green) and ECM components (magenta) on an MRC-5 cell. ECM components (1: fibronectin, 2: collagen type I, 3: laminin) were reconstituted as dSTORM movies.

**Supplemental Video 4.** Enlarged movie of the simultaneous observation of laminin by dSTORM (magenta) and a single sEV-CD63Halo7-TMR particle (green) on a living MRC-5 cell membrane. The field of view in this video is different from that of video 3.
